# Evolution with private resources reverses some changes from long-term evolution with public resources

**DOI:** 10.1101/2021.07.11.451942

**Authors:** Katrina van Raay, Sergey Stolyar, Jordana Sevigny, Adamaris Muñiz Tirado, Jeremy A. Draghi, Richard E. Lenski, Christopher J. Marx, Benjamin Kerr, Luis Zaman

## Abstract

A population under selection to improve one trait may evolve a sub-optimal state for another trait due to tradeoffs and other evolutionary constraints. How this evolution affects the capacity of a population to adapt when conditions change to favor the second trait is an open question. We investigated this question using isolates from a lineage spanning 60,000 generations of the Long-Term Evolution Experiment (LTEE) with *Escherichia coli*, where cells have access to a shared pool of resources, and have evolved increased competitive ability and a concomitant reduction in numerical yield. Using media-in oil emulsions we shifted the focus of selection to numerical yield, where cells grew in isolated patches with private resources. We found that the time spent evolving under shared resources did not affect the ability to re-evolve toward higher numerical yield. The evolution of numerical yield commonly occurred through mutations in the phosphoenolpyruvate phosphotransferase system. These mutants exhibit slower uptake of glucose, making them poorer competitors for public resources, and produce smaller cells that release less carbon as overflow metabolites. Our results demonstrate that mutations that were not part of adaptation under one selective regime may enable access to ancestral phenotypes when selection changes to favor evolutionary reversion.

## Introduction

The history of life has been inextricably shaped by evolutionary constraints. Given phenotypic trade-offs, an organism cannot simultaneously optimize two traits with respect to fitness (Agrawal et al. 2010; Stearns 1989). Thus, a population under strong selection to improve one trait may evolve a sub-optimal state for another trait. More generally, pleiotropy means that mutations in one gene affect multiple traits simultaneously (Pavličev & Cheverud, 2015), and antagonistic pleiotropy occurs when mutations are beneficial for one trait but detrimental for another. While there is a rich literature on evolutionary tradeoffs (Beardmore et al. 2011; Gounand et al. 2016; Guillaume & Otto, 2012) and antagonistic pleiotropy (Rose, 1982; Rose, 1985), there has been less attention given to how the evolutionary history of a lineage under selection for one trait affects its subsequent trajectory when selection shifts to favor a second trait that trades off with the first (but see Ostrowski et al. 2015; Teotónio & Rose, 2000; Travisano & Lenski, 1996; Velicer, 1999).

To understand how previous adaptation might affect future evolution, let us imagine a population under long-term selection for one trait that trades off with a second trait (Figure 1a). For simplicity, we assume higher values of each trait correspond to higher fitness when under selection. Long-term improvement in the favored trait (trait 1 in Figure 1a) occurs along with concomitant decreases in the second trait (trait 2). If selection shifted to favoring trait 2, how would the duration of history evolving under selection for trait 1 affect future evolution (Figure 1b)? There are at least two plausible hypotheses about how that duration would influence the rate of adaptation. The “Mutational Access” hypothesis (Figure 1c) posits that populations that spent more time under selection for trait 1 have fewer available mutations that would improve fitness by increasing trait 2 (i.e., the “late descendant” compared to the “intermediate descendant” in Figure 1a). If so, then this would lead to a *lower* rate of adaptation for the late descendant than for the intermediate one when trait 2 is favored. This mutational limitation could result from the necessity of compensating for (or reverting) more mutational steps; alternatively, it might occur if mutations increasing trait 2 become increasingly deleterious over time due to the entrenchment of trait 1 (Schank & Wimsatt, 2011; Shah et al. 2015). The “Strength of Selection” hypothesis (Figure 1d), by contrast, suggests that the intermediate and late descendants have equal access to mutations improving trait 2, but the selective benefits for the late descendant are greater due to the relatively larger changes in fitness upon improvement (Barrick et al. 2010; Chou et al. 2011). Under this hypothesis, the rate of adaptation is predicted to be *higher* for the late descendant.

**Figure 1:**
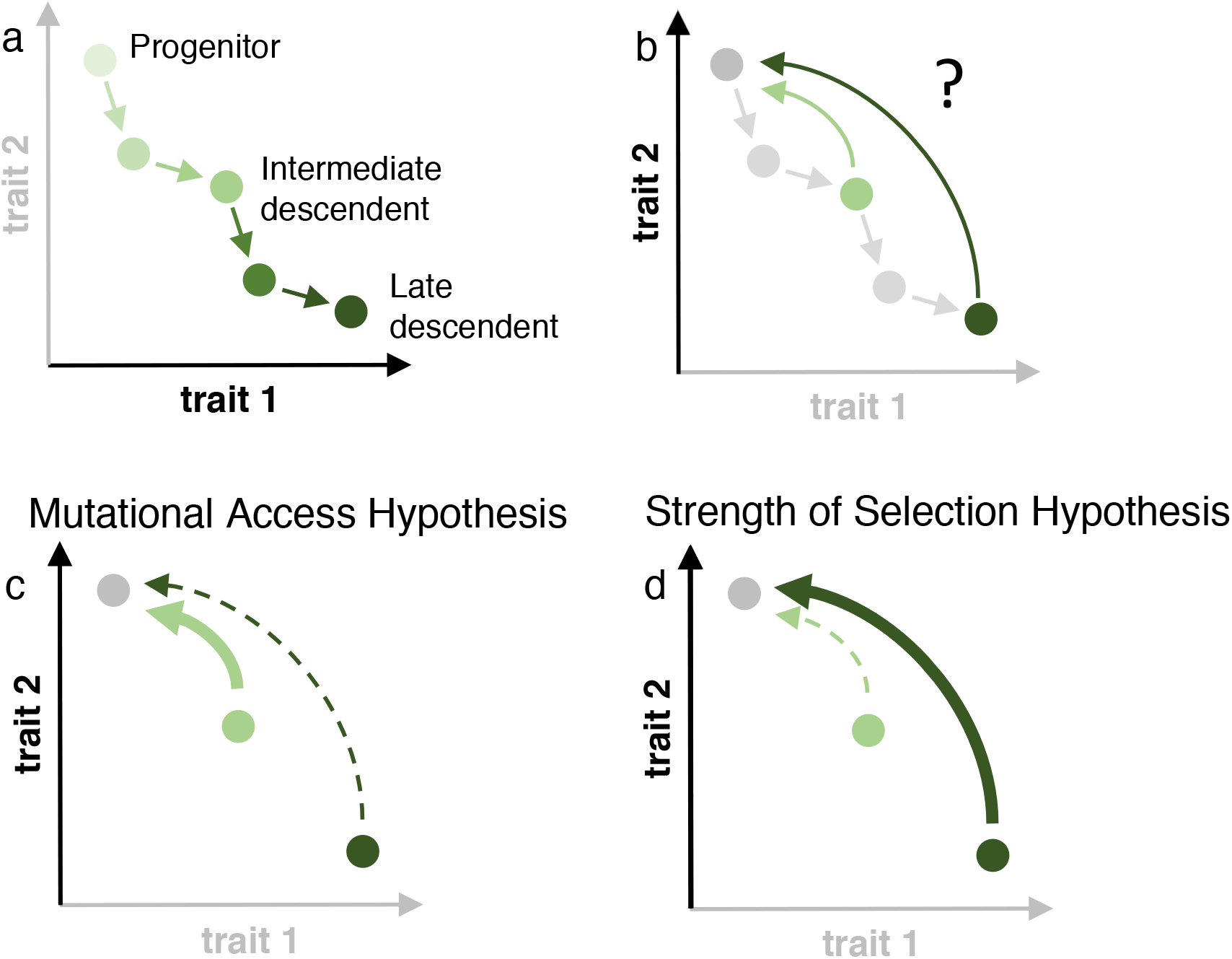
Evolutionary trajectories and alternative hypotheses. a) A population evolves under direct selection for one trait (trait 1, bolded axis). A higher value along either axis indicates an improvement in that trait under selection. Each circle represents a genotype in an evolutionary line of descent. We focus on three genotypes along an evolutionary trajectory: the progenitor, an intermediate descendent, and a late descendant. b) Selection is relaxed for trait 1 and strengthened for trait 2 (bolded axis). The curved arrow represents the transition of a population with descendent traits towards traits that resemble the progenitor when selection acts on trait 2. Whether, how fast, and by what mechanisms this transition occurs are the main questions addressed in this paper. c) Mutational access hypothesis. Intermediate (light green) and late (dark green) descendants are shown after selection is relaxed on trait 1 and strengthened on trait 2. The width of arrows pointing from each descendent towards the progenitor indicates the rate of change. The thick arrow from the intermediate descendent denotes a faster return to the progenitor phenotype than the dashed arrow from the late descendent. Under this hypothesis, the late descendent is either mutationally further from the progenitor state or the mutational options to revert to the progenitor phenotype are more costly due to entrenchment. d) Strength of selection hypothesis. Intermediate (light green) and late (dark green) descendants are shown when selection flips from favoring trait 1 to trait 2. The width of arrows pointing from each descendent towards the progenitor indicates the relative strength of selection. The thick arrow from the late descendent indicates stronger selection favoring a mutation that restores the ancestral phenotype, and thus a faster response, than for the same mutation in the intermediate descendant (dashed arrow). Under this hypothesis, both descendants have the same mutational access to various positions in phenotype space.

If adaptation is possible under selection for trait 2, then we expect a return towards the progenitor’s value of that trait. Does this phenotypic return occur by “rediscovering” the ancestral genetic and mechanistic underpinnings of the second trait (Levin et al. 1997; Pennings et al. 2020)? This rediscovery could occur by reverting key mutations, or by otherwise returning to the regulatory, physiological, or metabolic states that enabled the progenitor’s phenotype. Indeed, the mechanisms that underly a return to the ancestral state (e.g., Figures 1b-d) might differ depending on whether evolution is initiated with intermediate or late descendants.

Here we present a realization of the thought experiment in Figure 1, in which we test how the length of time that populations of *Escherichia coli* had been selected for faster growth affects their future adaptation in a novel environment that favors increased numerical yield (i.e., number of cells per concentration of substrate added). To do so, we take advantage of the Long-Term Evolution Experiment (LTEE), in which 12 replicate populations of *E. coli* have been evolving for over 70,000 generations in a relatively simple environment (Lenski 2017; Vasi et al. 1994; Wiser et al. 2013). Periodic samples from each line have been frozen in suspended animation, so that the “progenitor,” “intermediate descendant,” and “late descendant” are available by reviving samples from this living fossil record. The well-stirred, unstructured environment maintains a single, public pool of resources for the population and thus favors rapid growth, above all else (Vasi et al. 1994, Novak et al. 2006). Furthermore, there is no expectation of direct selection for either numerical yield or metabolic yield (proportion of carbon in the substrate that is incorporated into biomass). In an unstructured environment, a mutant that consumes resources more slowly but efficiently is disadvantaged, because a faster-growing but less-efficient competitor can deplete the shared resources in the meantime. Instead, a mutant that grows faster has an advantage in an unstructured environment, even if it leads to a reduction in the population’s numerical yield, metabolic yield, or both.

The outcomes in the LTEE are slightly more complicated, and they are mediated in part by changes to a third trait: cell size (Mongold & Lenski 1996). All of the LTEE lineages show increases in competitive ability relative to their common ancestor, while simultaneously having lower numerical yields (Vasi et al. 1994, Wiser et al. 2013). The reductions in numerical yield result from the evolution of much larger individual cells (Lenski & Travisano 1994, Vasi et al. 1994, Grant et al. 2021). In many bacteria, increased cell size is a physiological response to increased growth rate (Johnston et al. 1979; Pierucci, 1978). In addition, some mutations in bacteria affect both growth and cell size or shape (Monds et al. 2014, Yulo & Hendrickson 2019). Therefore, it is reasonable to expect that selection for faster growth has led to larger cells and, as a pleiotropic consequence, decreased numerical yield.

These evolutionary changes also extend to metabolic yield—the efficiency with which carbon sources are converted into biomass. During exponential growth on glucose, when most of the doublings occur during the LTEE daily cycle, direct measurements of metabolic yield through central metabolism show a small (but significant) decrease in the evolved populations. The excretion of acetate, an overflow metabolite, also increased by about 50% in the LTEE (Harcombe et al. 2013). Similar to the case with cell size, some loci with beneficial mutations in the LTEE lineages are known to affect acetate metabolism (Quandt et al. 2015); it is also known that rapid growth *per se* leads to increased overflow metabolite production in *E. coli* (Barrick & Lenski 2009; Farmer & Jones 1976). This situation is further complicated, however, by the finding that the LTEE populations have evolved increased total biomass when measured after a full 24-hour growth cycle (Lenski & Mongold 2000, Novak et al. 2006). While the increase might be caused in part by a change in biomass composition, it also reflects an increased ability to use excreted metabolites after the glucose has been depleted. Indeed, the evolved LTEE strains have increased their ability to grow on acetate, as well as to grow on other overflow metabolites that the LTEE ancestor could not use (Leiby & Marx, 2014). Although there was no direct advantage to increased biomass in a well-mixed environment, nonetheless biomass increased alongside faster growth in the LTEE. This constellation of changes can be understood by considering that, in the novel environment of the LTEE, the ancestral strain was likely far from the trade-off front between growth rate and total biomass, leaving room for improvements in both trats (Novak et al. 2006).

Now consider what might happen if the bacteria that previously evolved in the LTEE moved to and evolved in a structured environment that was otherwise similar to the unstructured LTEE environment. By distributing a transplanted population into many isolated ‘patches,’ each seeded by a single cell, any resources saved by growing more efficiently would accrue disproportionately to cells with the same genotype, which would effectively eliminate resource competition. Now imagine that the cells in these patches were periodically pooled, diluted distributed as single founders over a new set of empty patches. In that case, selection would favor increased numerical yield since each individual cell is a potential propagule able to establish a new subpopulation. Increased numerical yield could be achieved by increasing the metabolic yield, by producing smaller individual cells, or perhaps by some combination of these changes. In this way, selection shifts to favor numerical yield over competitive ability for shared resources (accomplishing the transformation from Figure 1a to Figure 1b in our thought experiment).

In our study, we achieve this shift by propagating cells in water-in-oil emulsions comprised of millions of aqueous media-filled droplets surrounded by an oil phase (Figure 2) (Bachmann et al. 2013). If a population of cells is diluted sufficiently before creation of the emulsion, then each cell will usually be the sole occupant of a droplet. In such a case, barring mutation, there is no competition between genotypes for resources, as each cell (and its progeny) has access to a private resource supply demarcated by the droplet. If growth of isolated cells inside droplets proceeds for sufficient time to exhaust the substrate and is followed by demulsification, dilution, and redistribution into droplets across successive transfers (as in Figure 2a), then those genotypes that produce *more* cells (rather than the fastest-growing cells) will have an advantage. Indeed, prior experiments have demonstrated that various forms of isolation can favor numerical yield at the expense of competitive ability in several different biological systems (Bachmann et al. 2013; Eshelman et al. 2010; Kerr et al. 2006; van Tatenhove-Pel et al. 2021).

**Figure 2:**
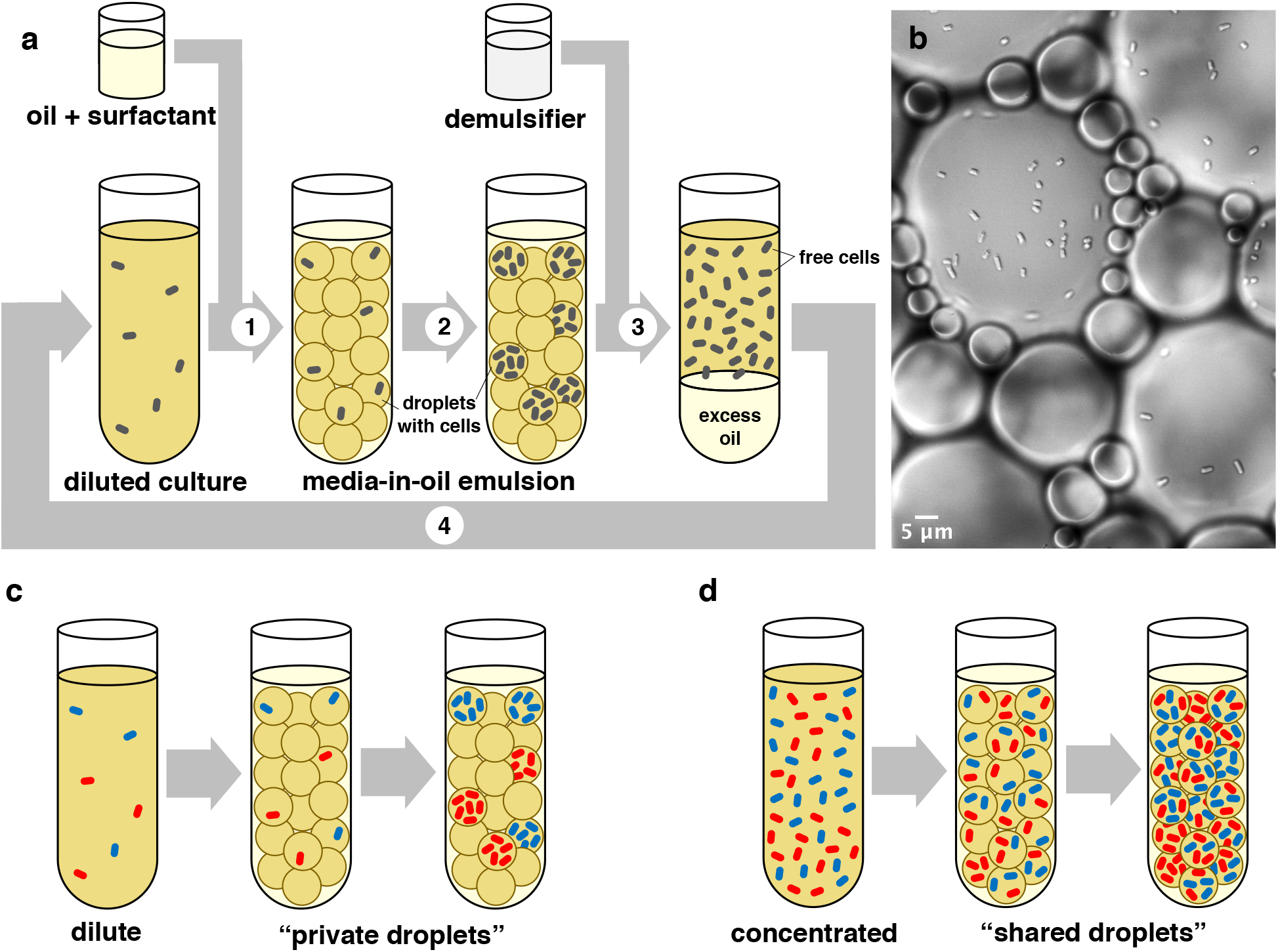
**a)** Schematic of transfer protocol: 1. A mixture of oil and surfactant is added to a diluted culture of bacteria in growth media and vortexed to form millions of droplets, 2. Bacteria in the emulsion are incubated to allow subpopulation growth within the droplets, 3. After overnight growth, a demulsifier is added to break the emulsion and resuspend the bacteria, 4. The free cells are diluted into fresh media and the cycle is repeated, **b)** Micrograph of bacteria growing in emulsion droplets, **c)** Schematic of inoculation and growth for private droplets, **d)** Schematic of inoculation and growth for shared droplets. For parts c and d, the two colors of bacterial cells represent distinct genotypes.

Here, we instantiate the thought experiment of Figure 1 by considering two bacterial traits: competitive ability (the relative increase in proportion of a focal strain when competing with another strain for shared resources) and numerical yield (the absolute number of cells produced by a focal strain when resources are not shared). We focus on a single LTEE lineage in order to explore how previous selection for one trait (competitive ability) affects the tempo and mode of evolution under selection for a second trait (numerical yield) in the presence of a phenotypic tradeoff. We probe how the length of time adapting to the well-mixed LTEE regime affects the ability of a population to adapt under the new emulsion condition. We examine whether increases in numerical yield during growth with private resources come with associated decreases in competitive ability for shared resources, and whether they are achieved through changes in metabolic yield and associated overflow metabolite production, decreases in cell size, or by both factors. Finally, by determining the nature and number of mutations that occurred, we can investigate whether any evolutionary reversions to the ancestral phenotypes were caused by changes in the same loci/processes that were implicated in the evolution of fast growth in the original LTEE environment.

## Methods

### Strains

All experiments were founded with *E. coli* B isolates that came from the Ara–5 lineage in the LTEE. The isolates used in our experiment came from 0 generations (the founder of the LTEE), 20,000 generations, and 60,000 generations of evolution in the LTEE (Table S1; hereafter called 0K, 20K, and 60K, respectively). The 0K isolate is the ultimate progenitor of the 20K and 60K isolates in the context of the LTEE. However, because we conduct evolution experiments in the emulsion system with these three strains, we will refer to these LTEE isolates as the “0K emulsion ancestor”, the “20K emulsion ancestor”, and the “60K emulsion ancestor”.

### Emulsion evolution experiment

Following the pioneering work of Bachmann et al. (2013), we propagated three replicate cultures founded by each of the three LTEE isolates (the emulsion ancestors) under selection for increased numerical yield by using water-in-oil emulsions for 100 transfers. This method (see below) creates millions of media-filled, picoliter-sized droplets surrounded by an oil phase, in which bacterial cells in one droplet are isolated from those in others (Figure 2). Emulsions were set up in 1.5-ml Eppendorf conical microcentrifuge tubes, and they were incubated at 37°C without shaking.

In order to select for increased numerical yield, we diluted the bacterial culture before setting up the emulsions, such that 90% of occupied droplets would contain only 1 cell at the beginning of each transfer cycle. We subsequently showed that occupied droplets had an average of 1.12 cells (see Supplementary section *Calculating Droplet Statistics*). Bacteria were allowed to grow and divide in droplets for 24 hours, after which the droplets were broken with 1*H*,1*H*,2*H*,2*H*-perfluoro-1-octanol, releasing all the cells into a common pool. A fraction of the cells from this pool were immediately used to initiate the next transfer (see Figure 2a). Because of the level of dilution before emulsification, lineages from this treatment have evolved in “private droplets” (see Figure 2c). This treatment was continued for 100 transfers.

To control for any effects of growing in emulsions, we also propagated three replicate cultures founded by each LTEE isolate for 100 transfers under higher starting-density conditions, in which each occupied droplet had almost 3 cells on average (see Supplementary section *Calculating Droplet Statistics*). We predicted that we would not observe evolution towards increased numerical yield in this “shared-droplet” treatment (see Figure 2d).

During the evolution experiment, the cell density of freshly broken emulsions was measured with a FilterMax F5 Multi-Mode Microplate Reader (Molecular Devices) after 24 hours. Briefly, we pipetted 150 μl of overnight culture from a broken emulsion into a flat-bottom 96-well microtiter plate, and we recorded an OD_595_ reading. This reading was then used to estimate cell density using a calibration curve giving the relationship between OD_595_ and CFU/ml, which was established in prior experiments. Cultures were then diluted to 2 × 10^6^ CFU/ml for the private droplet treatment and 5 × 10^7^ CFU/ml for the shared droplet treatment. Emulsions were created by adding 200 μl of a mixture of 9 parts Novec 7500 HFE to 1 part Pico-surf (5% (w/w)) to 300 μl of diluted culture in the growth medium DM1000 in 1.5 ml Eppendorf tubes, and vortexing at maximum speed for 2 minutes. DM1000 contains 1000 mg/l of glucose, and it was used instead of the DM25 (25 mg/l glucose) used in the LTEE because DM1000 exhibited clearer differences between the 0K and 60K samples in terms of numerical yield and competitive ability than DM25 in pilot experiments.

Single-colony isolates were picked from each population at the end of our evolution experiment (after transfer 100) and frozen at −80 °C as 20% v/v glycerol stocks. We refer to these evolved isolates as “emulsion descendants.” Two loci previously shown to distinguish the 0K, 20K, and 60K ancestors were Sanger sequenced for each isolate to confirm the intended derivation of the isolates. We detected within-experiment contamination via Sanger and subsequent whole-genome sequencing in 4 of the 18 emulsion descendants, so these populations were dropped from our analysis. The complete genomes of the remaining 14 strains (8 private droplet, 6 shared droplet) were sequenced and analyzed, and the 6 isolates of private-droplet descendants that had mutations were used for further analysis. For the shared droplet treatment, one descendent was randomly chosen from each lineage to be assessed in the phenotypic assays (3 isolates).

### Competition assays

Using the frozen samples of the emulsion ancestors (0K, 20K, and 60K) and their emulsion descendants from the private and shared droplet treatments, we assessed the relative competitive abilities of all of our strains. The emulsion descendants competed against a common marked strain (equivalent to our 0K emulsion ancestor) in shared droplet conditions. We used a neutral marker (a point mutation in the *araA* gene) to distinguish the focal and common competitor strains on tetrazolium-arabinose indicator agar plates.

Competitions ran for 3 transfer cycles in emulsions. Competitions were initiated with a 1:1 ratio of the focal strain (denoted *F*) to the common competitor (denoted *C*). The competitive ability of the focal strain relative to the common competitor (*w*(*F*, *C*)) was assessed by calculating a ratio of their realized Malthusian growth rates (Lenski et al. 1991):

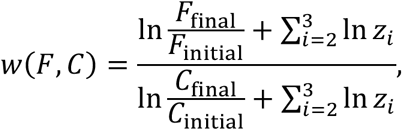

where *X*_initial_ and *X*_final_ are the initial (beginning of the first transfer) and final (end of the third transfer) densities, respectively, of strain *X* (with *X* ∈ {*F*, *C*}) and *z*_*i*_ is the factor by which the population is diluted in order to initiate the *i*^th^ transfer.

### Genome Sequencing

Single-colony isolates from 14 evolved lines and their 3 ancestors were grown in DM1000 and frozen as 20% v/v glycerol stocks at −80 °C. For genome sequencing, cells from these frozen stocks were reanimated in 5 ml of LB (lysogeny broth). Genomic DNA was extracted using a Qiagen DNEasy kit with 1 ml of overnight culture, following the manufacture’s protocol. Library preparation and sequencing for 16 strains was done at the Microbial Genome Sequencing Center (MiGS) at the University of Pittsburgh. One strain (the 0K emulsion ancestor) was sequenced at the University of Washington on Illumina’s NextSeq platform and prepped with Illumina Nextera barcodes. Genome sequences had an average of 131X coverage.

Mutations (predicted mutations, unassigned missing coverage, and unassigned new junctions) were called using breseq (Version 0.33.2) (Deatherage & Barrick, 2014) with default parameters. All mutations were confirmed using Integrative Genomics Viewer (Robinson et al. 2011), and four point mutations identified in strains that differed phenotypically from their ancestors in cell size or numerical yield were Sanger sequenced for confirmation (Table S4).

### Numerical yield and cell size assays

Populations of each emulsion ancestor and descendant were grown under the shared-droplet conditions over a standard 24-hour growth cycle. We used the shared-droplet emulsions instead of the well-stirred conditions to control for the microenvironmental effects of growing in an emulsion (e.g., oxygen depletion during growth). The final cell number in the 24-hour sample was used as our measure of numerical yield. Cell size was measured as the median cell volume (μm^3^) of several thousand cells using a Beckman Coulter MSE 4 instrument.

### Metabolic analysis

Supernatants from the yield assays were filtered and stored at −80 °C prior to metabolic analysis. After thawing, samples were analyzed using High Performance Liquid Chromatography (HPLC) on a 20A chromatographic system (Shimadzu) equipped with diode array and refractive index detectors. Samples were eluted with 6 mM H2SO4 at a flow rate of 0.6 ml/min from an Aminex HPX 87H organic acid analysis column (300 by 7.8 mm) (BioRad). We recorded the areas for all noticeable peaks associated with detected metabolites (glucose, citrate, acetate, formate, and lactate), and these values were converted to metabolite concentrations (in mM) using organic acid and sugar standards (from BioRad or made in-house). Citrate, an iron chelator present in DM medium, is neither produced nor consumed by these bacterial strains, and it has been shown not to affect the growth of the Ara–5 lineage (Leiby et al. 2012). Thus, citrate was used as an internal control.

### Analysis of metabolic data

The rates of production for each metabolite are difficult to compare because of differences in the growth rates of the strains. To minimize this potentially confounding factor, we performed linear regression of the concentration of each metabolite against the remaining concentration of glucose. The magnitude of the slope of this relationship estimates the ratio of metabolite produced for each unit of glucose consumed, assuming an approximate equilibrium between the uptake of glucose and the release of metabolites into the medium. As glucose levels drop, however, there is increased uncertainty in the glucose measurements; moreover, as the glucose becomes scarce, the cells may begin to consume the excreted metabolites, which would bias our estimates of the production rates. To avoid these problems, we excluded time-points in the linear regressions for which the mean glucose concentration was below 0.8 mM. We chose this threshold because, at lower concentrations, growth curves showed clear signs of departure from exponential growth. We also excluded the measurements made at t=0 and, instead, we constrained the x-intercept of the regressions to reflect the known initial concentrations in the media (i.e., 6 mM glucose and none of the overflow metabolites). To estimate how quickly the glucose was consumed, we performed regressions of glucose concentrations against time, using the same criteria for data exclusion. This approach provided a measure of growth rate that was insensitive to differences in cell size across strains.

## Results & Discussion

### Private-droplet treatment selected for high numerical yield

Our private droplet treatment was designed to minimize local competition between genotypes, thereby focusing selection upon increased numerical yield. To assess whether the emulsion-evolved populations underwent changes in numerical yield, we measured their cell densities, along with those of their ancestors, at the end of 24 hours of growth in the emulsion system. Providing some reassurance that salient features of the LTEE environment are maintained in our emulsion cultures, the decreases in numerical yield seen during the LTEE were also detected in emulsion -- the 20K and 60K ancestors had decreased numerical yield compared to the 0K ancestor (Figure 3, Table S3; p < 0.01, unpaired one-tailed t-tests). All 6 private droplet descendants had significantly increased numerical yield (cell number) compared with their direct ancestor (Figure 3, Table S2; BK 433: p < 0.05; BK 431, BK 437, BK 438, BK 441, BK 442: p < 0.01, unpaired two-tailed t-tests). Our shared-droplet treatment served as control for the private-droplet treatment, allowing local competition between genotypes, while controlling for growth in the emulsion environment. Two of the three shared-droplet descendants had increased numerical yield and one had decreased yield; however, the gains were less than those in the private-droplet treatment, and all three changes were non-significant. Combining the 0K, 20K, and 60K lines, the private-droplet descendants were significantly more productive than the shared-droplet descendants (p=0.039, paired one-tailed t-test; see Supplementary section *Statistical Analysis*). The two evolutionary treatments had different effective population sizes. However, the weaker numerical-yield response in the shared-droplet treatment relative to the private-droplet treatment cannot be explained by the lack of mutational opportunity, because the number of cell divisions per transfer is actually greater in the shared-droplet treatment (see Supplementary section *Calculating Number of Cell Divisions in Droplets*). Taken together, these data confirm our prediction that the private-droplet treatment exerts stronger selection for increased numerical yield than does the shared-droplet treatment.

**Figure 3:**
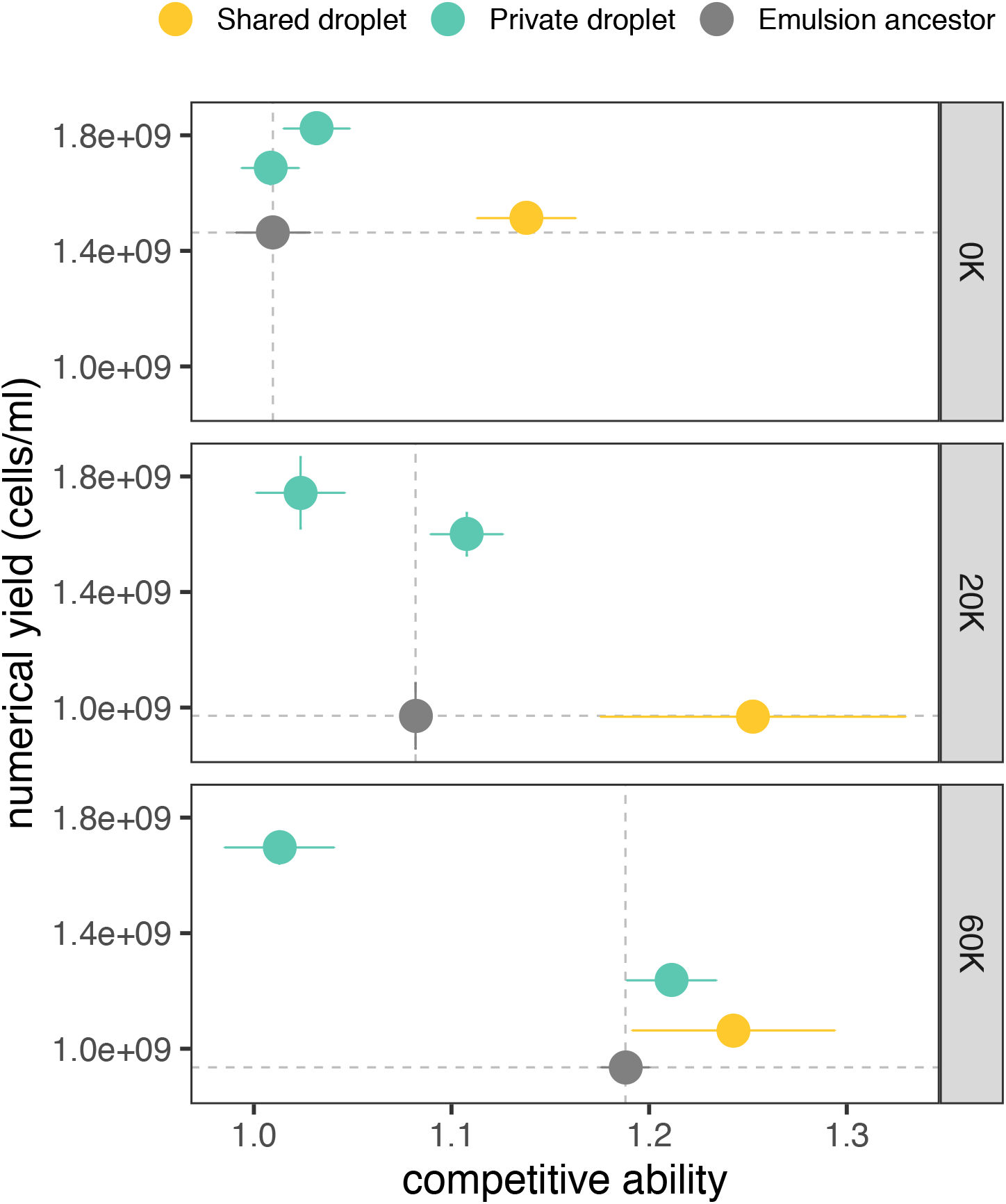
Numerical yield (cell density after the standard 24-hour growth cycle) compared to relative competitive ability, both measured in the emulsion system. Each panel shows strains originating from a single emulsion ancestor (in grey, at right) from the Ara–5 lineage of the LTEE (0, 20,000, and 60,000 generations). Colors indicate emulsion evolution treatments (labels at top). The coordinate positions of each point are the averages of three replicate assays for each phenotypic trait. Error bars show SEM; when they are not visible along either axis, the corresponding SEM is smaller than the symbol itself. Dashed lines show the mean competitive ability and numerical yield of the relevant ancestor.

In particular, the shared-droplet results demonstrate that the increase in numerical yield in the private-droplet treatment was not simply an evolutionary response to propagation under the emulsion conditions. Instead, these results imply that the initial density within a droplet affects the evolutionary outcome. By reducing competition between genotypes within the private droplets, selection has favored numerical yield.

### Shared droplet treatment favored competitive ability

When resources are shared, selection for greater competitive ability is expected. To assess competitive ability, a focal strain was paired with a marked version of the 0K ancestor and proportions were tracked under high density emulsion conditions such that resources were shared between competitors. Again, emulsion populations showed consistent results with the batch culture environment used during the LTEE -- the competitive ability of the 20K ancestor was elevated compared to the 0K ancestor, and the 60K ancestor had a further increase over the 20K ancestor (Figure 3,Table S3; p < 0.05 unpaired, one-tailed t-tests). Competitive ability increased in all three shared-droplet descendants that we tested (Figure 3). However, the change was significant relative to the corresponding emulsion ancestor only in the case of the line derived from the 0K ancestor (Table S2). Competitive ability also increased in three of the six private-droplet descendants, but none of these increases were significant, and the magnitudes of the increases were smaller than those observed in the shared-droplet treatment. Competitive ability decreased in the other three private-droplet descendants, and the decline was significant in one case (BK 441: p < 0.01, unpaired two-tailed t-test). Combining the results from the 0K, 20K, and 60K lines, the shared-droplet descendants were significantly more competitive than the private-droplet descendants (p = 0.038, paired one-tailed t-test; see Supplementary section *Statistical Analysis*). Overall, these results support our prediction that starting each droplet with multiple cells favors the evolution of increased competitive ability.

### Tradeoff between competitive ability and numerical yield

If competitive ability and numerical yield strictly traded off, then competitive ability would decrease in the private-droplet treatment as numerical yield increased, and numerical yield would decrease in the shared-droplet treatment as competitive ability increased. However, we did not see such a strict tradeoff. Although the private-droplet descendants had larger gains in numerical yield, some shared-droplet descendants also experienced moderate gains in numerical yield. Similarly, although the shared-droplet descendants exhibited larger gains in competitive ability, some private-droplet descendants also showed moderate improvements in their competitive ability. Thus, these two traits do not strictly tradeoff with one another, and instead there seems to be some misalignment, in which larger gains in one trait sometimes occur with more modest gains in the other trait.

To develop this notion of misalignment further, we can contrast the case of a strict tradeoff, or pure antagonistic pleiotropy (Figure 4a), with two different scenarios of synergistic pleiotropy. The first synergistic scenario involves strict alignment, such that a mutation that is most beneficial for trait 1 is also the most beneficial for trait 2 (Figure 4b). In essence, this scenario is the opposite of strict antagonistic pleiotropy. The second form of synergism involves misalignment. In this scenario, all mutations simultaneously improve both traits, but the mutations that are the strongest for trait 1 are the weakest for trait 2, and *vice versa* (Figure 4c). Misalignment occurs when the relative effect of a mutation on one trait is the opposite of its relative effect on another trait. Therefore, antagonistic pleiotropy (Figure 4a) is also a case of misalignment, although the term is broader than the traditional definition of “antagonism” (as illustrated in Figure 4c). In the Supplementary section, we develop a statistical test for misalignment. Using this test, we find that the increases in numerical yield in the private-droplet treatment came with concomitant weaker performance in competitive ability, while the improvements in competitive ability in the shared-droplet treatment came with weaker performance in numerical yield. Despite the absence of a strict tradeoff, these traits exhibited misalignment in our experiment. When two traits are misaligned, independent evolution in environments that favor one trait or the other can produce a pattern that supports a canonical tradeoff curve when the descendants are pooled across environments and the ancestors are ignored (i.e., considering only the colored points, while ignoring the grey points, in Figures 3 and 4c).

**Figure 4:**
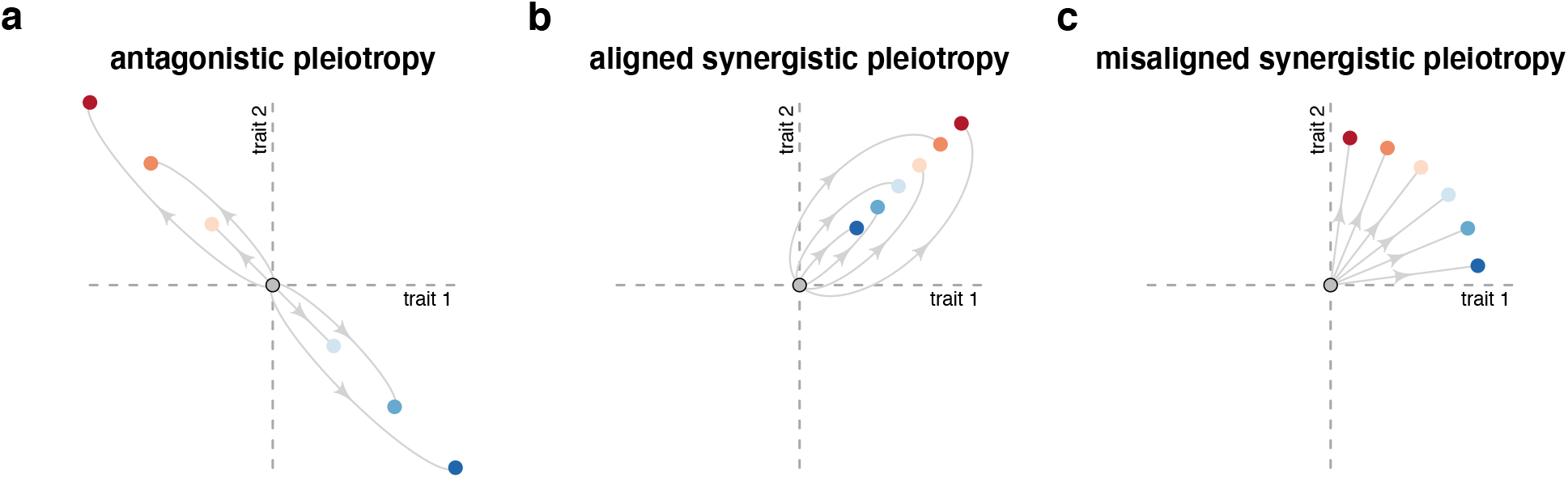
Three scenarios for pleiotropy with respect to two traits. The grey circle is the focal, ancestral genotype and the colored circles show the phenotypes of evolved genotypes. Here, fitness improvement with regard to a trait is taken to mean a *positive* change relative to the ancestor. (a) Antagonistic pleiotropy under a “strict tradeoff” situation, in which an improvement in one trait leads to a decline in the other, and the magnitude of enhancement in one dimension correlates with the magnitude of deterioration in the other. (b) Aligned synergistic pleiotropy describes a situation where the magnitude of improvement in one trait correlates with the magnitude of improvement in the other. (c) Misaligned synergistic pleiotropy describes a situation in which evolution improves both traits, but those genotypes with the strongest improvement in one trait have the weakest improvement in the other.

### Private-droplet treatment favored smaller cells

While performing microscopy to assess droplet diameter (Supplemental Figures S8 & S9), we noticed variation in cell size across the emulsion treatments. The cells that had evolved in private droplets appeared smaller than cells that had evolved in shared droplets. We therefore decided to investigate this trait systematically. Figure 5 shows the relationship between cell size and numerical yield.

**Figure 5:**
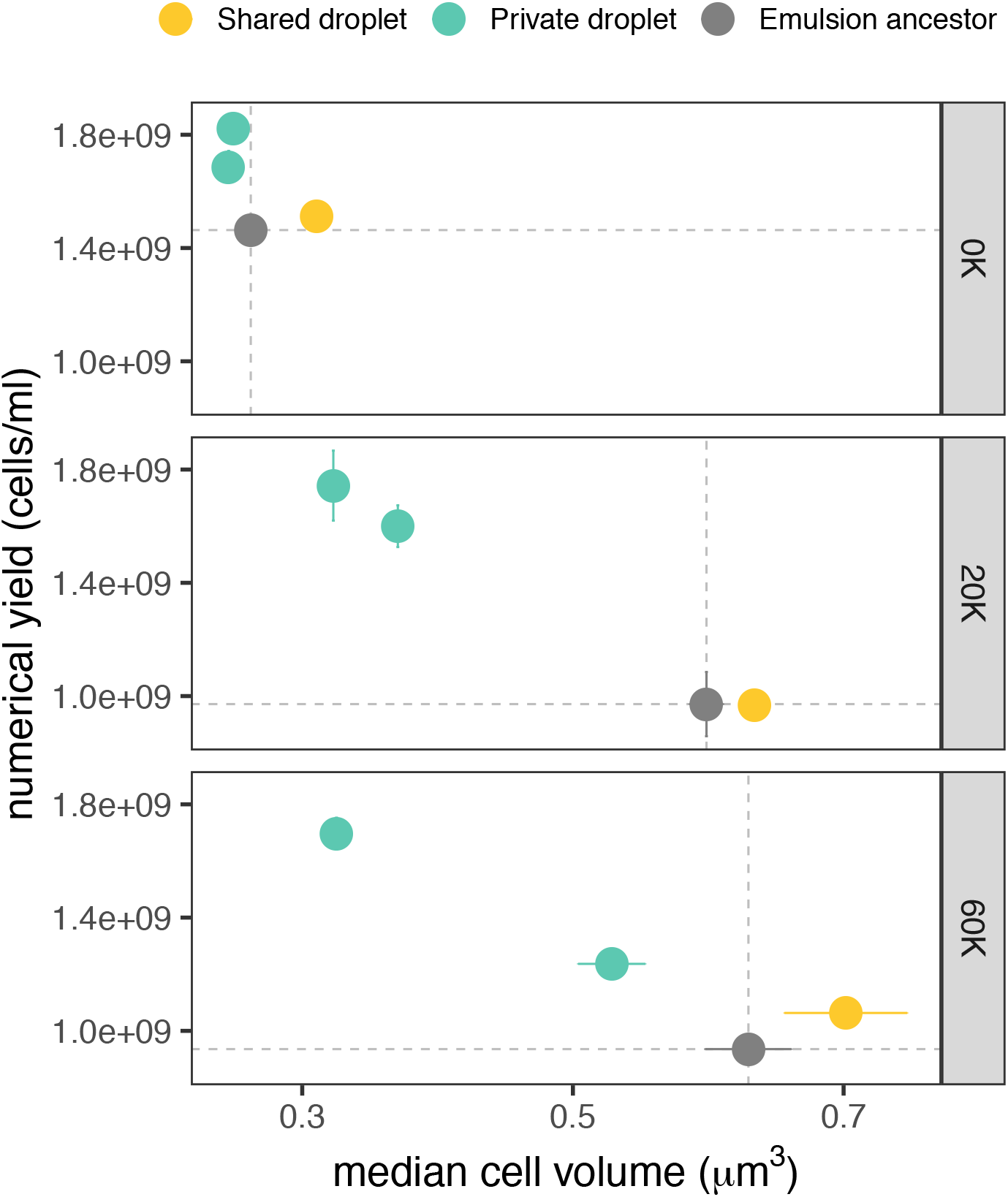
Median cell volume compared to numerical yield. Each panel shows strains originating from a single emulsion ancestor (in grey, at right) from the Ara–5 lineage of the LTEE (0, 20,000, and 60,000 generations). Colors indicate emulsion evolution treatments (labels at top). The coordinate positions of each point show the average of three replicate assays for each phenotypic trait. Error bars show SEM; when they are not visible along either axis, the corresponding SEM is smaller than the symbol itself. Dashed lines show the mean numerical yield and cell volume of the relevant ancestor.

Both cell size and total biomass increased in all 12 LTEE populations during selection for increased competitive ability for shared resources despite the fact that neither is directly a target of selection under those conditions (Grant et al. 2021, Lenski & Mongold, 2000). This increase in cell size can be seen in our data for the three time points that we sampled from the Ara–5 population, by comparing the values of the three grey points along the x-axis in Figure 5. For the emulsion evolved lines, the median cell volume decreased in all of the isolates from the private-droplet treatment, and the changes were significant in three cases (BK 437, BK 438, BK 441: p < 0.01, unpaired, two-tailed t-test). Using an emulsion protocol similar to our private-droplet treatment, Bachman *et al*. (2013) also reported observing an evolved decrease in cell size in *Lactococcus lactis*. They noted that propagation in an emulsion system could favor such a change because any mutant that distributed the same total amount of biomass into a greater number of smaller cells would be able to colonize more droplets each transfer. Our shared-droplet treatment provides an appropriate control for this inference. Indeed, the median cell size actually increased slightly in all three replicates of this treatment, and the change was significant in one case (BK 443: p < 0.01, unpaired, two-tailed t-test). The opposing directional changes in our two treatments indicates that neither the emulsion environment nor the transfer protocol (pooling all droplets and transferring only a fraction of cells to the next growth cycle, as shown in Figure 2a) was responsible for the consistently smaller cells seen in the private-droplet descendants. Instead, these findings support the hypothesis that smaller cells evolved in response to the reduced inter-genotype competition in the private-droplet treatment. When the 0K, 20K, and 60K lines are pooled, the difference in cell size between the private-droplet and shared-droplet descendants is marginally significant (p=0.051, paired one-tailed t-test; see Supplementary section *Statistical Analysis).*

### Genome sequencing reveals mutations in phosphoenolpyruvate phosphotransferase system

Given the evolution of smaller cells in the private-droplet treatment, we wondered if these populations were reverting to the ancestral state of the LTEE. After all, the progenitor of the LTEE (the 0K ancestor in our emulsion experiment) had very small cells. In particular, would the same genes and metabolic pathways that changed in the LTEE and led to larger cells in that experiment undergo reversion? Or would novel genetic and metabolic changes lead to a similar phenotypic endpoint? Also, would these changes depend on the particular LTEE-derived ancestor used in the emulsion experiment (e.g., 20K versus 60K)? To find out, we sequenced the complete genomes of 14 emulsion-evolved descendants and their three corresponding ancestors.

This whole-genome sequencing revealed a total number of 13 mutations that distinguish the evolved descendants from their ancestors (Table S4). We saw no cases of identical mutations in multiple descendants, nor was any gene mutated in descendants that evolved in the two different treatments. However, three of the six descendants from the private-droplet treatment had point mutations in genes encoding proteins in the phosphoenolpyruvate (PEP) phosphotransferase system (PTS) (Table 1), which is involved in glucose uptake (Carmona et al. 2015; Escalante et al. 2012; Nam et al. 2001; New et al. 2014; Notley-Mcrobb et al. 2006; Xia et al. 2017). All three of these descendants also had increased numerical yield and reduced cell size (Table S2), and all three were derived from the 20K and 60K emulsion ancestors. Of the three private-droplet descendants without mutations in PTS genes, two derived from the 0K ancestor, which already had high numerical yield in the emulsion conditions. The third private-droplet descendant without mutations in PTS genes derived from the 60K ancestor, and it exhibited much smaller changes in numerical yield and cell size than its counterpart with a PTS-associated mutation. The changes in numerical yield and cell size were also small for both of the private-droplet descendants derived from the 0K ancestor, suggesting that the high-yield phenotype of the LTEE progenitor was largely maintained. Bachmann et al. (2013) also found a point mutation in a PTS gene in one strain that evolved increased numerical yield in emulsions, suggesting that PTS might have some role in modulating numerical yield.

**Table 1:** Mutations in private-droplet descendants associated with PTS.

| STRAIN (BK#) | LTEE GEN | POSITION | REF | ALT | ANNOTATION | GENE | DESCRIPTION |
| --- | --- | --- | --- | --- | --- | --- | --- |
| 437 | 20K | 2462747 | A | C | *86Y (TAA→TAC) | *ptsH* | phosphocarrier protein Hpr |
| 437 | 20K | 3414187 | G | A | E55K (GAA→AAA) | *crp* | cAMP-activated global transcriptional regulator CRP |
| 438 | 20K | 1173797 | G | A | G454S (GGT→AGT) | *ptsG* | PTS glucose transporter subunit IIBC |
| 441 | 60K | 2462629 | T | C | L47P (CTG→CCG) | *ptsH* | phosphocarrier protein Hpr |

The PTS-associated mutations occurred in *ptsH*, which encodes a phosphocarrier protein; *ptsG*, which encodes a subunit of the glucose transporter; and *crp*, which encodes a global transcriptional regulator that affects some steps in glucose uptake (Kimata et al. 1997). Given that these mutations include an early stop codon and non-conservative changes in amino acids, it is likely that some or all of them cause reductions or even losses of function. Mutations in *pykF*, which encodes pyruvate kinase, were among the earliest and most repeatable genetic changes in the LTEE (Woods et al. 2006; Tenaillon et al. 2016; Peng et al. 2018). These mutations presumably reduce the conversion of PEP to pyruvate in central metabolism, which would leave more PEP available to power the transport of PTS-dependent sugars. Indeed, distinct patterns of pleiotropy were observed in the LTEE for various substrates that differed in terms of whether their uptake requires the PTS (Travisano & Lenski, 1996). Thus, although the mutations selected in the private droplet-treatment were not in the same genes as those responsible for phenotypic changes in the LTEE, they appear to affect the same system.

### Slower glucose uptake and decreased overflow metabolism in private-droplet descendants

Perhaps the private-droplet descendants achieved their higher numerical yield and smaller cell size in a physiologically similar way to the LTEE progenitor, despite genetic changes that did not merely reverse those that occurred during the LTEE. Because the mutations we found in the PTS system would be predicted to affect glucose transport, we decided to directly examine whether the private-droplet descendants had evolved toward a metabolic state, at least in terms of glucose use and overflow metabolite production, that resembled the LTEE progenitor.

We examined the concentrations of both glucose and various excreted metabolites in spent medium as the various bacterial strains grew in emulsions (Figure 6a). We observed that the glucose concentration for the 20K and 60K private-droplet descendants decreased more slowly than for the corresponding shared-droplet descendants, their ancestors, and the 0K descendants of either the shared or private droplets (Figure 6a, glucose panel). The slower glucose consumption was most pronounced in the private-droplet descendants that had mutations in PTS, indicating that these mutations likely simultaneously reduce the rate of glucose utilization while increasing numerical yield.

**Figure 6:**
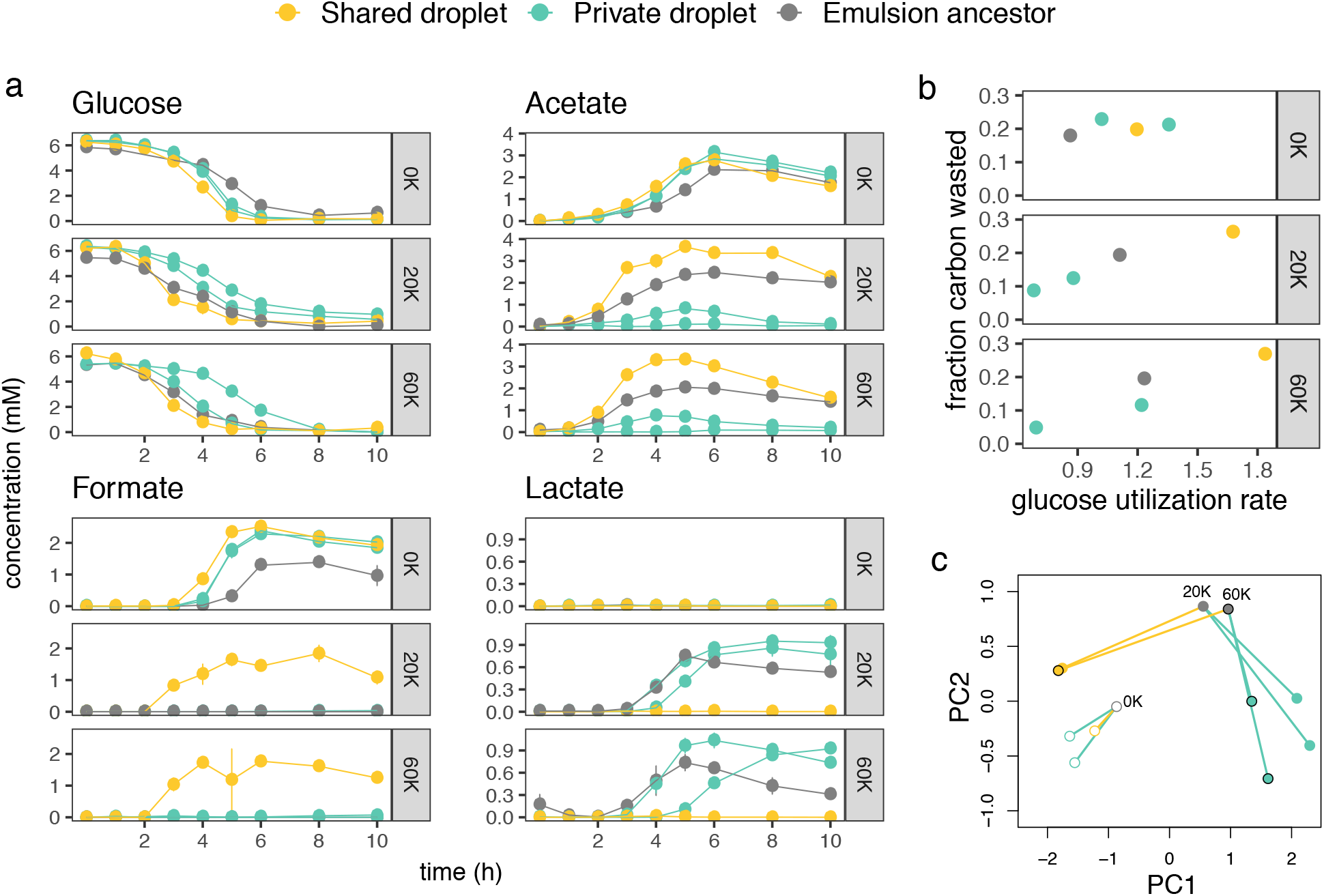
Glucose consumption and metabolite production. a) The concentrations of glucose, acetate, formate and lactate over 10 hours. Each trajectory shows one strain, and each point along a trajectory is the average of up to three replicates per strain. Colors indicate emulsion evolution treatments (labels at top). b) Glucose utilization versus the fraction of carbon wasted. c) Principal components analysis (PCA) based on the strain-specific rates of excreted metabolites per unit glucose consumed. The 0K strains are shown as empty points, the 20K strains as filled points without borders, and the 60K strains as filled points with borders. Lines, colored by treatment, connect emulsion ancestors (grey points) to their descendants.

The primary metabolite that is excreted by *E. coli* growing on glucose, and by the LTEE lines in particular, is acetate (Harcombe et al. 2013). Acetate is a byproduct of both glycolysis and fermentation, which is a less efficient form of metabolism than respiration (Szenk et al. 2017). The 0K emulsion evolution decedents did not markedly differ in metabolic signatures from the LTEE ancestor. However, we saw differences in acetate production between the private-droplet descendants from 20K and 60K and their emulsion ancestors. Specifically, the private-droplet descendants produced less acetate, which suggests they were more efficient at glucose metabolism under these conditions than their ancestor. In contrast, some of the shared-droplet descendants accumulated higher levels of acetate (Fig. 6a). In both cases, the acetate concentration eventually declines as it is consumed by the cells after they have depleted the available glucose. Interestingly, in emulsion-evolved populations of a different strain of *E. coli* (MG1655), Rabbers et al. (2021) found an increased numerical yield was achieved with no change in acetate production. Those results are consistent with our observations for the 0K descendants but differ from what we observe in the 20K and 60K descendants, which suggests a history of rate adaptation may leave its mark on future evolution.

In addition to acetate, other organic acids such as formate and lactate can be generated by E. *coli* during growth on glucose. For all of the isolates in our study, either formate or lactate was generated, but not both. For the 0K lineages, formate was produced by all types (ancestor, private-droplet descendants, and shared-droplet descendant). For the 20K and 60K lineages, only the shared-droplet descendant produced formate, whereas lactate was produced by the 20K and 60K ancestors and their private-droplet descendants.

Differences between strains in the accumulated concentrations of excreted metabolites might reflect different production rates, but they might also be influenced by differences in growth rate, as well as subsequent consumption of those products. We therefore standardized these measurements by computing the rate of production of each excreted metabolite per unit of glucose consumed, and by excluding time points after the remaining glucose fell below a threshold concentration (see *Methods—Analysis of metabolic data*). After so doing, Figure 6b shows a positive relationship between the initial rate of glucose utilization and the fraction of this carbon that is “wasted” as excreted metabolites in the 20K and 60K emulsion lines. Carbon waste was calculated as a sum of the concentrations across all excreted metabolites concentrations, weighted by the number of carbon atoms in each metabolite. The data indicate that the shared-droplet descendants use the glucose rapidly, but in a relatively wasteful way, whereas the private-droplet descendants consume glucose more slowly and efficiently.

We used these standardized rates (Table S5) to perform a principal components analysis (PCA; Figure 6c) in order to visualize the differences between the emulsion ancestors and their descendants. The metabolic changes between the 0K ancestor and its descendants were modest, but the changes in the 20K and 60K lines were much greater. For the 20K and 60K ancestors, the metabolic shifts in the private-droplet treatment were distinct from those in the shared-droplet treatment, while the overall patterns of change were nearly identical for the 20K and 60K lines. Thus, in addition to PTS-related mutations conferring high numerical yield for three of the four private-droplet descendants of these two ancestors, the magnitude and nature of the metabolic changes appear similar. One private-droplet descendant (derived from the 60K ancestor) exhibited a smaller decrease in cell size and a more moderate increase in numerical yield (Figure 5). This isolate also did not have mutations associated with PTS, and it differed somewhat less in its metabolic profile from its ancestor. On balance, therefore, we lack clear support for either the Mutational Access Hypothesis or the Strength of Selection Hypothesis (Figure 1). Nonetheless, it is clear that just one or a few novel genetic changes can restore, at least approximately, the ancestral phenotypes of slow growth, small cells, and high numerical yield, even after tens of thousands of generations that led to faster growth, larger cells, and lower numerical yield.

### Summary

Three of the four private-droplet descendants of the 20K and 60K ancestors achieved or even surpassed the numerical yield of the ultimate LTEE progenitor (the 0K emulsion ancestor). When compared to their ancestors, these high-yield descendants showed several parallel changes: smaller cells, more efficient glucose metabolism, mutations in genes encoding the PTS, and lower levels of acetate production. None of these changes occurred in the descendants that evolved from the same ancestors under the shared-droplet treatment. While the increased numerical yield and smaller cell size appeared to be reversions to the phenotypic state of the ultimate progenitor, the genetic and metabolic underpinnings of the descendants’ phenotypes were distinct from those of the ultimate progenitor. Specifically, mutations in genes encoding the PTS were not responsible for evolutionary improvement in growth rate during the LTEE, and the metabolic profiles of the 20K and 60K private-droplet descendants were distinct from that of the 0K emulsion ancestor (Fig. 6c). Therefore, these data indicate a novel mechanistic basis for phenotypic reversion in this system. Specifically, mutations in the PTS likely involved a reduction in glucose uptake, thereby lowering the growth rate (and leading to a reduced competitive ability). However, these same mutations appear to lead to both smaller size and limited loss of carbon via overflow metabolites, thereby improving numerical yield. Notably, the same kind of mutations produced the phenotypic reversion for both the 20K and 60K emulsion lines.

Dollo’s law of irreversibility states that “An organism never returns exactly to a former state, even if it finds itself placed in conditions of existence identical to those in which it has previously lived. But by virtue of the indestructibility of the past [...] it always keeps some trace of the intermediate stages through which it has passed” (Gould, 1970). Our experiment certainly did not return these organisms to their ancestral conditions. However, the private-droplet treatment was designed to place populations under selection for an ancestral phenotype, namely higher yield. Even as the private-droplet lineages derived from later generations of the LTEE recovered the high numerical yield of the LTEE ancestor, their phenotypic “return” did not involve a straightforward reversal of the steps in their evolutionary history. Rather, the mutational targets and metabolic patterns underlying their return were distinct from those evolutionary paths explored in the LTEE. By contrast, when under the same selection for increased numerical yield, the private-droplet descendants of the 0K emulsion ancestor exhibited genetic and metabolic changes distinct from the 20K and 60K emulsion lines, suggesting that changes during the first 20,000 generations of the LTEE had altered the potential for subsequent adaptation to the private-droplet regime Such contingency is a recurring feature of adaptation in biological systems (Blount et al. 2008; Card et al. 2019). However, we found no evidence of contingency in the later generations, as both the intermediate (20K) and late (60K) isolates from the LTEE readily recovered their ancestral form, seemingly by taking similar steps. Therefore, past evolution need not invariably alter future evolution, even when the future involves a return to the past.

