## Supplemental file for "Evolution with private resources reverses some changes from long-term evolution with public resources"

### A return to form: Novel mutations keep ancestral traits accessible despite evolutionary history

**Page Contents**

| Page | Contents |
|---|---|
| 2 | **Supplemental Experimental Methods** |
| 2 | *Microscopy for droplet size* |
| 2 | Figure S1 |
| 2 | *Calculating droplet statistics* |
| 5 | Figure S2 |
| 7 | *Calculating number of cell divisions in droplets* |
| 9 | Figure S3 |
| 10 | *Calibrating OD readings and cell counts* |
| 10 | *Optical density and changes in cell size* |
| 11 | *Focal strains* |
| 11 | *Growth and cell size assays* |
| 12 | Figure S4 |
| 13 | **Statistical and Quantitative Methods** |
| 13 | *Statistical analysis* |
| 13 | *Misalignment analysis* |
| 15 | Figure S5 |
| 16 | Figure S6 |
| 17 | Figure S7 |
| 18 | Figure S8 |
| 19 | **List of Strains and Supplemental Results** |
| 19 | Table S1 |
| 20 | Table S2 |
| 21 | Table S3 |
| 22 | Table S4 |

#### Supplemental Experimental Methods

##### *Microscopy for droplet size*

Emulsion droplets were imaged using an inverted water-immersion Olympus confocal microscope. Images were made by pipetting 40 µl of emulsion onto a glass slide with a glass coverslip on the bottom and part of the slide carved out to make a cavity. Images were analyzed in FIJI (version 2.0.0-rc-69/1.52n), and the maximum width of 238 droplets was measured (in µm) using the line tool.

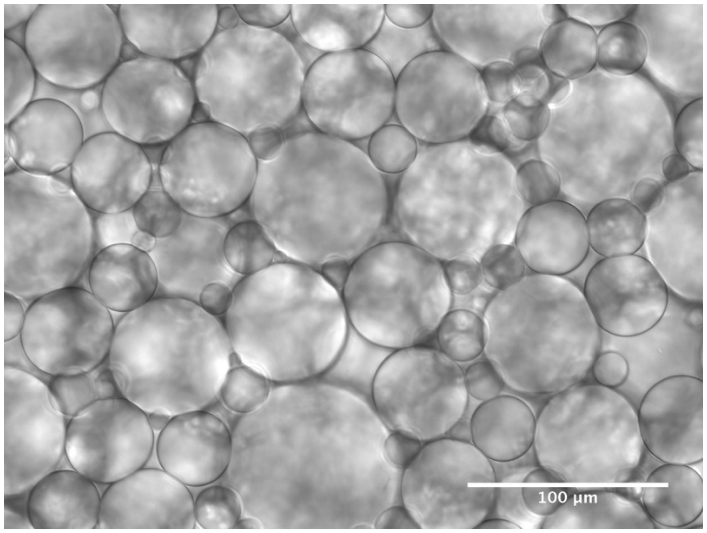


**Figure S1:** An example image of emulsion droplets using confocal microscopy.

##### *Calculating droplet statistics*

Droplets in the emulsions were not uniform in size (see Fig. S1). We can use the distribution of droplet volumes to estimate how many cells, on average, were in an occupied droplet for each of our emulsion treatments (shared or private).

We can describe the distribution of droplet volumes with a probability density function $p(v)$. Suppose that the density of cells before emulsification is given by $\lambda$ (CFU/ml). The probability that a droplet has $c$ cells, $\delta(c)$, is:

$$\delta(c) = \int_0^\infty \frac{e^{-\lambda v}(\lambda v)^c}{c!} p(v)dv.$$

The mean number of cells per droplet would then be:

$$\mu = \sum_{c=0}^{\infty} c\delta(c).$$

This summation can be reframed as follows:

$$\mu = \sum_{c=1}^{\infty} \int_0^\infty \frac{e^{-\lambda v}(\lambda v)^c}{(c-1)!} p(v)dv,$$

$$\mu = \int_0^\infty p(v)dv \left\{ \sum_{c=1}^{\infty} \frac{e^{-\lambda v}(\lambda v)^c}{(c-1)!} \right\},$$

$$\mu = \int_0^\infty p(v)dv \left\{ \lambda v \sum_{c=1}^{\infty} \frac{e^{-\lambda v}(\lambda v)^{c-1}}{(c-1)!} \right\},$$

$$\mu = \int_0^\infty p(v)dv \left\{ \lambda v \sum_{i=0}^{\infty} \frac{e^{-\lambda v}(\lambda v)^i}{i!} \right\},$$

$$\mu = \int_0^\infty \lambda v p(v)dv,$$

$$\mu = \lambda \bar{v},$$

where $\bar{v}$ is the average volume of a droplet.

If we condition on the presence of at least one cell in the droplet, we can recompute the mean number of cells, which we call $\mu_{\text{occupied}}$. In that case, we obtain:

$$\mu_{\text{occupied}} = \frac{\sum_{c=1}^{\infty} c\delta(c)}{\sum_{c=1}^{\infty} \delta(c)},$$

$$\mu_{\text{occupied}} = \frac{\mu}{\int_0^{\infty} p(v)dv \left\{\sum_{c=1}^{\infty} \frac{e^{-\lambda v}(\lambda v)^c}{c!}\right\}},$$

$$\mu_{\text{occupied}} = \frac{\mu}{\int_0^{\infty} p(v)dv \left\{-e^{-\lambda v} + \sum_{c=0}^{\infty} \frac{e^{-\lambda v}(\lambda v)^c}{c!}\right\}},$$

$$\mu_{\text{occupied}} = \frac{\mu}{\int_0^{\infty} p(v)dv \{1 - e^{-\lambda v}\}},$$

$$\mu_{\text{occupied}} = \frac{\int_0^{\infty} \lambda v p(v)dv}{1 - \int_0^{\infty} e^{-\lambda v} p(v)dv}.$$

Finally, the fraction of occupied droplets that have only a single cell, $\rho_{\text{single}}$, is given by:

$$\rho_{\text{single}} = 1 - \frac{\sum_{c=2}^{\infty} \delta(c)}{\sum_{c=1}^{\infty} \delta(c)},$$

$$\rho_{\text{single}} = 1 - \frac{\int_0^{\infty} p(v)dv \left\{\sum_{c=2}^{\infty} \frac{e^{-\lambda v}(\lambda v)^c}{c!}\right\}}{1 - \int_0^{\infty} e^{-\lambda v} p(v)dv},$$

$$\rho_{\text{single}} = 1 - \frac{\int_0^{\infty} p(v)dv \left\{-e^{-\lambda v} - e^{-\lambda v}\lambda v + \sum_{c=0}^{\infty} \frac{e^{-\lambda v}(\lambda v)^c}{c!}\right\}}{1 - \int_0^{\infty} e^{-\lambda v} p(v)dv},$$

$$\rho_{\text{single}} = 1 - \frac{\int_0^{\infty} p(v)dv \{1 - e^{-\lambda v} - e^{-\lambda v}\lambda v\}}{1 - \int_0^{\infty} e^{-\lambda v} p(v)dv},$$

$$\rho_{\text{single}} = 1 - \frac{1 - \int_0^{\infty} e^{-\lambda v} p(v)dv - \int_0^{\infty} e^{-\lambda v}\lambda v p(v)dv}{1 - \int_0^{\infty} e^{-\lambda v} p(v)dv},$$

$$\rho_{\text{single}} = \frac{1 - \int_0^{\infty} e^{-\lambda v} p(v)dv - 1 + \int_0^{\infty} e^{-\lambda v} p(v)dv + \int_0^{\infty} e^{-\lambda v}\lambda v p(v)dv}{1 - \int_0^{\infty} e^{-\lambda v} p(v)dv},$$

$$\rho_{\text{single}} = \frac{\int_0^\infty e^{-\lambda v} \lambda v p(v) dv}{1 - \int_0^\infty e^{-\lambda v} p(v) dv}.$$

If we can determine the values of the following integrals

$$\mu = \int_0^\infty \lambda v p(v) dv,$$

$$\alpha = \int_0^\infty e^{-\lambda v} p(v) dv,$$

$$\beta = \int_0^\infty e^{-\lambda v} \lambda v p(v) dv,$$

then we can compute $\mu_{\text{occupied}} = \frac{\mu}{1-\alpha}$ and $\rho_{\text{single}} = \frac{\beta}{1-\alpha}$.

We have an empirically determined distribution of droplet diameters, measured in μm. Figure S2 shows a distribution of droplet diameters measured on 238 droplets, with the droplets binned in 10-μm intervals. Suppose we consider one of these intervals, ranging from diameter $D_{\text{low}}$ to $D_{\text{high}}$. Letting the volume of a droplet with diameter $D$ be given by $V(D)$, the average droplet volume in the focal interval will be:

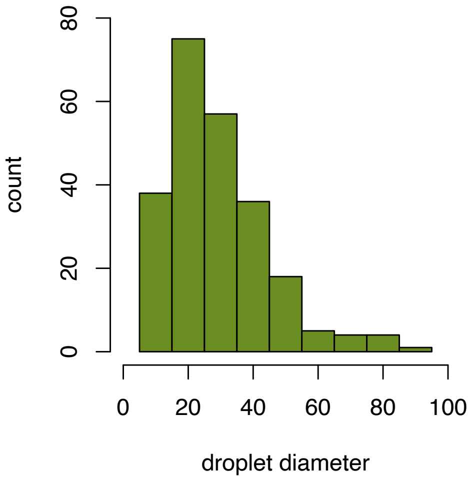

**Figure S2**: The distribution of droplet diameters from microscopic analysis.

$$V(D_{\text{low}}, D_{\text{high}}) = \frac{\int_{D_{\text{low}}}^{D_{\text{high}}} V(D) dD}{D_{\text{high}} - D_{\text{low}}}.$$

Because $V(D) = \frac{\pi}{6} D^3$,

$$V(D_{\text{low}}, D_{\text{high}}) = \left(\frac{\pi}{6}\right) \frac{\int_{D_{\text{low}}}^{D_{\text{high}}} D^3 dD}{D_{\text{high}} - D_{\text{low}}},$$

$$V(D_{\text{low}}, D_{\text{high}}) = \left(\frac{\pi}{6}\right)\frac{\frac{D_{\text{high}}{}^4}{4} - \frac{D_{\text{low}}{}^4}{4}}{D_{\text{high}} - D_{\text{low}}},$$

$$V(D_{\text{low}}, D_{\text{high}}) = \left(\frac{\pi}{24}\right)\frac{\left(D_{\text{high}}{}^2 - D_{\text{low}}{}^2\right)\left(D_{\text{high}}{}^2 + D_{\text{low}}{}^2\right)}{D_{\text{high}} - D_{\text{low}}},$$

$$V(D_{\text{low}}, D_{\text{high}}) = \left(\frac{\pi}{24}\right)\frac{\left(D_{\text{high}} - D_{\text{low}}\right)\left(D_{\text{high}} + D_{\text{low}}\right)\left(D_{\text{high}}{}^2 + D_{\text{low}}{}^2\right)}{D_{\text{high}} - D_{\text{low}}},$$

$$V(D_{\text{low}}, D_{\text{high}}) = \left(\frac{\pi}{24}\right)\left(D_{\text{high}} + D_{\text{low}}\right)\left(D_{\text{high}}{}^2 + D_{\text{low}}{}^2\right),$$

If the droplet count for our focal bin is $n$ and the total number of droplets is $N$, then we will say the probability of falling in the interval is

$$P(D_{\text{low}}, D_{\text{high}}) = \frac{n}{N}$$

So, let us designate the diameter boundaries of the bins as $D_0$, $D_1$, $D_2$, $D_3$, … $D_{\text{B}}$. We can then use the following approximations:

$$\mu = \int_0^\infty \lambda v p(v) dv \approx \sum_{i=0}^{\text{B}-1} \lambda\{V(D_i, D_{i+1})\}\{P(D_i, D_{i+1})\},$$

$$\alpha = \int_0^\infty e^{-\lambda v} p(v) dv \approx \sum_{i=0}^{\text{B}-1} e^{-\lambda\{V(D_i, D_{i+1})\}}\{P(D_i, D_{i+1})\},$$

$$\beta = \int_0^\infty e^{-\lambda v} \lambda v p(v) dv \approx \sum_{i=0}^{\text{B}-1} e^{-\lambda\{V(D_i, D_{i+1})\}} \lambda\{V(D_i, D_{i+1})\}\{P(D_i, D_{i+1})\}.$$

There is one remaining issue, namely the units. The standard units for cell density are CFU/ml, whereas the volume unit for droplets here is μm$^3$. However, because there are $10^{12}$ μm$^3$ per milliliter, we simply divide the cubic-micron volume by $10^{12}$ in the integrals above.

For the private-droplet treatment, the cell density was 2 x $10^6$ CFU/ml and the statistics are:

$$\mu = 0.05 \text{ cells/droplet}$$
$$\mu_{\text{occupied}} = 1.12 \text{ cells/droplet}$$
$$\rho_{\text{single}} = 0.90$$

For the shared-droplet treatment, the cell density was 5 x $10^7$ CFU/ml and the statistics are:

$$\mu = 1.33 \text{ cells/droplet}$$
$$\mu_{\text{occupied}} = 2.99 \text{ cells/droplet}$$
$$\rho_{\text{single}} = 0.45$$

Therefore, in the private-droplet treatment, 90% of all occupied droplets have only a single cell, with an average of 1.12 cells per occupied droplet. In the shared droplet treatment, by contrast, 45% of the occupied droplets have a single cell, with an average of almost 3 cells per occupied droplet.

***Calculating number of cell divisions in droplets***

Here, we focus on the number of cell divisions in the emulsion system if the initial cell density is $\lambda_i$ (CFU/ml). Suppose also that, at stationary phase, any droplet that was initialized with one or more cells will reach a final density of $\lambda_f$ (where $\lambda_f > \lambda_i$). Focusing on one milliliter of volume, the initial number of cells is $c_i = \lambda_i$. The final number of cells ($c_f$) will not generally be $\lambda_f$, however, because some droplets are not occupied. The total number of cell divisions is simply $d = c_f - c_i$, because every cell produced requires a cell division. The number of generations, measured as population doublings, is then given by $g = \log_2(c_f/c_i)$.

So, to find the number of cell divisions, we focus on finding $c_f$. For a droplet with volume $v$ (measured in ml), the probability that it gets colonized by one or more cells (given the initial density $\lambda_i$) is

$$\chi(v) = 1 - e^{-\lambda_i v}$$

We assume that all colonized droplets reach a final density of $\lambda_f$. Further, we assume that every colonized droplet is initiated at a cell density less than $\lambda_f$, such that no cell death occurs before reaching the final density. These assumptions are reasonable for the initial densities and droplet-size distribution of our experiment. The final number of cells in a single milliliter is

$$c_f = \frac{\int_0^\infty \chi(v)\lambda_f v p(v)dv}{\int_0^\infty v p(v)dv}$$

Noting $\bar{v} = \int_0^\infty vp(v)dv$, we can simplify this expression

$$c_f = \frac{\lambda_f \int_0^\infty (1 - e^{-\lambda_i v}) v p(v) dv}{\bar{v}} = \frac{\lambda_f \bar{v} - \lambda_f \int_0^\infty e^{-\lambda_i v} v p(v) dv}{\bar{v}}$$

Letting $\beta = \int_0^\infty e^{-\lambda_i v}\lambda_i v p(v)dv$, we have:

$$c_f = \lambda_f - \frac{\lambda_f \beta}{\lambda_i \bar{v}}$$

Therefore, the number of cell divisions is

$$d = \lambda_f\left(1 - \frac{\beta}{\lambda_i \bar{v}}\right) - \lambda_i$$

Using terminology from the previous section, we obtain

$$\bar{v} = \int_0^\infty vp(v)dv \approx \sum_{i=0}^{\mathrm{B}-1}\{V(D_i, D_{i+1})\}\{P(D_i, D_{i+1})\}$$

$$\beta = \int_0^\infty e^{-\lambda_i v}\lambda_i v p(v) dv \approx \sum_{i=0}^{\mathrm{B}-1} e^{-\lambda_i\{V(D_i,D_{i+1})\}}\lambda_i\{V(D_i, D_{i+1})\}\{P(D_i, D_{i+1})\}$$

Using the droplet distribution from Figure S2, in Figure S3 we graph the logarithm (base 10) of the number of cell divisions for a range of initial cell densities (from $10^6$ to $10^8$ CFU/ml) and three final cell densities ($5\times10^8$, $1\times10^9$, and $2.5\times10^9$ CFU/ml). These final densities are in the range of the final density in our experimental system. As the initial cell density rises, two effects occur. First, the number of droplets that are colonized increases. Second, the number of cells per occupied droplet increases. The first factor leads to more cell divisions, whereas the second factor leads to fewer divisions. As the initial cell density is increased from very low values, the first effect dominates the second, and the number of cell divisions increases. However, as the initial cell density continues to increase towards the final cell density, the first effect becomes greatly muted (as nearly all the droplets are occupied) and the second effect dominates, so that the number of cell divisions decreases (this effect can be seen in the curve for the $5\times10^8$ final density). Figure S3 also shows that, for each of the final cell densities, the shared-droplet treatment has more cell divisions than the private-droplet treatment.

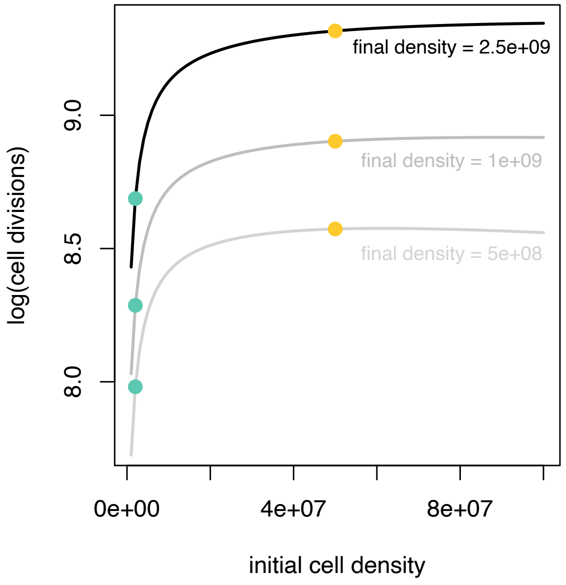


**Figure S3**: The logarithm of the number of cell divisions (per ml) as a function of initial cell density (CFU/ml) for three different final cell densities (CFU/ml). The private-droplet and shared-droplet treatments are defined by their initial densities, and they are indicated by teal and yellow points, respectively.

##### *Calibrating OD readings and cell counts*

In order to facilitate calculating the dilutions to perform in our experimental transfers, we built a JavaScript calculator that converted each $OD_{595}$ reading to a cell-density estimate. We used that estimate to calculate the appropriate dilutions and volume of media to add for each population. This tool is available online: https://kerrlab.github.io/Project_ET/

To calibrate the OD to CFU relationship, we initially performed three replicate mock competitions between REL606 and REL11638 in emulsions. We used this mixture to capture some of the variation in OD values between clones starting from different evolutionary timepoints in our experiments. After overnight growth, the emulsions were broken and a dilution series was prepared on a 96-well plate. The OD values of these dilutions were measured, and multiple dilutions were plated to obtain CFU counts. With known dilution series and CFUs, we had a large range of OD values and CFU counts. Using a second-degree polynomial fit to these data, the calculator estimated the CFU count from an entered OD value. The calculator was further refined after collecting additional data with corresponding CFU and OD values.

***Optical density and changes in cell size***

Optical density (OD) is a measurement of the amount of light absorbed by a sample. As cells in a culture multiply, the culture becomes more turbid and its optical density increases. Cell size can also affect OD. Generally, for the same density of cells, a culture of smaller cells will have lower OD (Stevenson et al. 2016). Because we used OD to compute the dilutions for transfers in our evolution experiment (see previous section), and because we did not measure cell size over the course of the experiment, evolutionary changes in cell size might inadvertently affect our dilution scheme. In the private-droplet treatment cells evolved smaller size, such that our conversion of OD to cell density would have led to an underestimate. This underestimation would have led to a higher initial cell density each transfer, and it would thereby have *reduced* the selection for numerical yield. Thus, our finding of increased numerical yield in the private-droplet treatment is conservative, having occurred despite the weakened selection. Similarly, the slight increases in cell size in the shared-droplet treatment would have led to a lower initial cell density each transfer, thereby reducing selection for competitive ability, and again making the inferences conservative.

***Focal strains***

We used six evolved isolates from the private-droplet treatment that had acquired one or more mutations during the experiment (as determined by whole-genome sequencing) in the cell size, numerical yield, and metabolic assays. We excluded two private-droplet isolates that did not acquire mutations or show changes in cell size or numerical yield from additional assays and analyses. For the shared-droplet treatment, we chose one isolate at random from the three replicates for each emulsion ancestor (0K, 20K and 60K) for further analysis.

***Growth and cell size assays***

To assess population growth, we revived frozen isolates in 5 ml of DM1000 and grew them overnight at 37 °C with shaking. We added 5 µl of each overnight culture to 10 ml of Isoton and ran samples through a 30-µm aperture on a MSE4 Beckman Coulter Counter to estimate cell density. The cultures were then diluted to 5 x $10^7$ cells/ml, and 30 emulsions were set up for each isolate to allow 3 replicates at 10 timepoints each. The emulsion populations were allowed to grow without shaking at 37 °C, and they were then destructively sampled at 0, 1, 2, 3, 4, 5, 6, 8, 10, and 24 hours. The cell density and median cell volume (µ$m^3$) were recorded at each timepoint using the Coulter Counter (Figure S4).

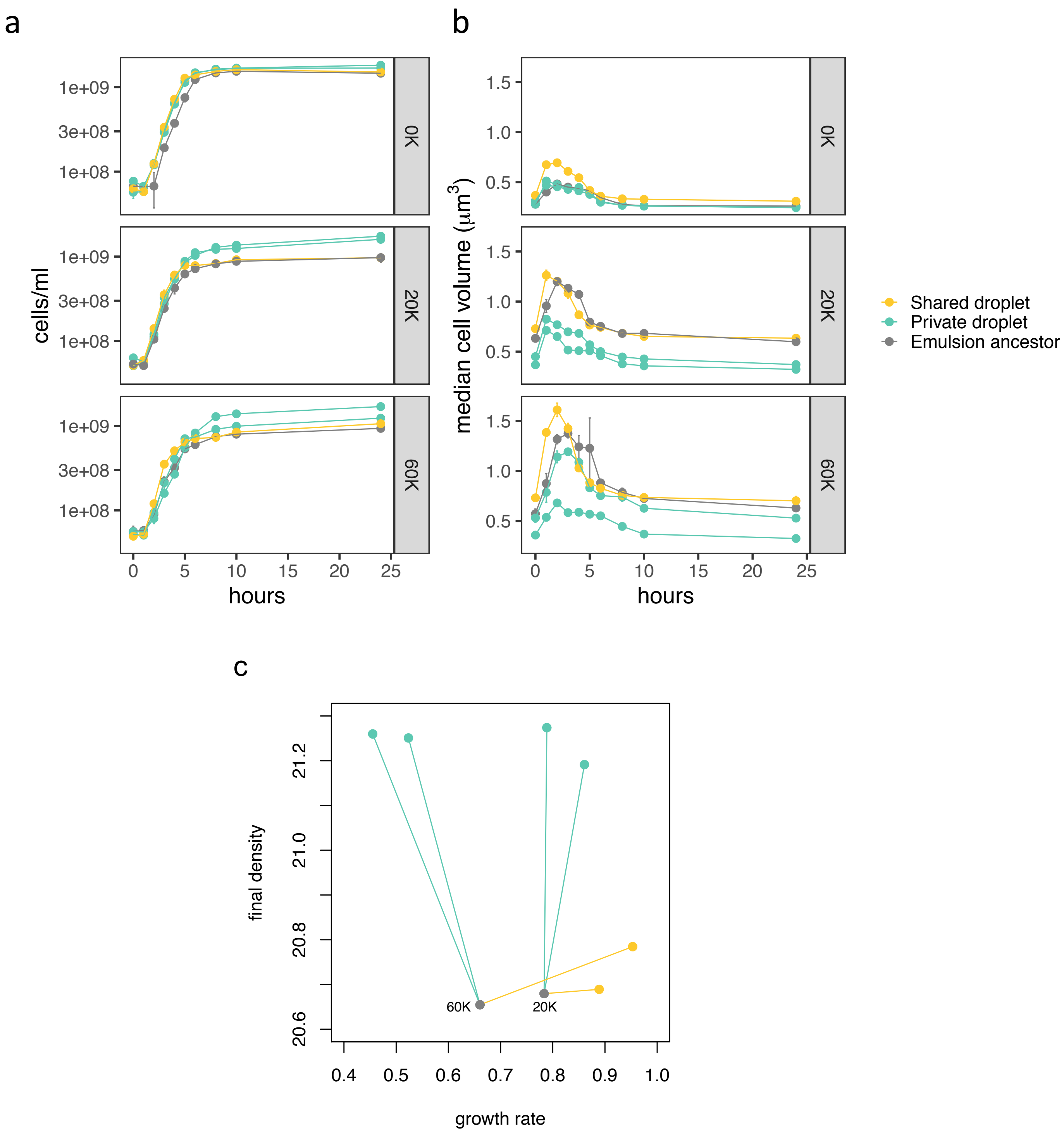


**Figure S4:** Population growth and median cell volume. **a)** Cell density over a 24-hour transfer cycle. **b)** Median cell volume over the same 24-hour cycle. In panels **a** and **b**, each curve shows one strain colored according to the labels (at right). Each point is the average of three replicates, and error bars are SEM. **c)** The growth rate ($hr^{-1}$) between 1 and 3 hours and the final density (log(CFUs/mL)) are shown for the 20K and 60K lines. The private droplet descendants have higher final density and lower growth relative to the matched shared droplet descendant.

#### Statistical and Quantitative Methods

##### *Statistical analysis*

All statistical analyses were performed using R (Version 3.6.0). Means and standard error of the mean were calculated for competitive fitness, cell size, and cell number for each isolate at each timepoint, usually based on three replicate samples. For each of the above traits, we ran unpaired, two-tailed t-tests comparing the emulsion-evolved strains with their emulsion ancestors (for 0K, 20K, and 60K lineages).

To compare the sets of descendants from the private-droplet and shared-droplet treatments, we used the mean values for each isolate across all technical replicates. We additionally took the average value for the two private-droplet lines derived from each emulsion ancestor (0K, 20K, and 60K). Pairing each private-droplet average with the corresponding shared-droplet value based on the common emulsion ancestor, we performed one-tailed, paired t-tests on the mean values of competitive fitness, cell size, and numerical yield. We used one-tailed tests here because our alternative hypotheses were directional.

##### *Misalignment analysis*

We consider a general case in which experimental evolution occurred in two treatments ($\boldsymbol{B}$ and $\boldsymbol{C}$) and we are tracking two phenotypic traits ($x$ and $y$) in multiple lines for each treatment. Treatment $\boldsymbol{B}$ selects for increased values of trait $x$, while treatment $\boldsymbol{C}$ selects for increased values of trait $y$.

We assume that the vector $\vec{a} = \langle x_a, y_a \rangle$ describes the phenotype of the common ancestor of all the evolved lines in both treatments. In line $\ell$ of the $\boldsymbol{B}$ treatment, the evolved phenotype is denoted

$\vec{b}_\ell = \langle x_{b_\ell}, y_{b_\ell} \rangle$. Similarly, in line $\ell$ of the $\boldsymbol{C}$ treatment, the evolved phenotype is denoted $\vec{c}_\ell = \langle x_{c_\ell}, y_{c_\ell} \rangle$. We start by rewriting all evolved phenotypes as deviations from their common ancestor. Specifically, line $\ell$ of the $\boldsymbol{B}$ and $\boldsymbol{C}$ treatments are expressed as:

$$\tilde{b}_\ell = \vec{b}_\ell - \vec{a} = \langle x_{b_\ell} - x_a, y_{b_\ell} - y_a \rangle = \langle \tilde{x}_{b_\ell}, \tilde{y}_{b_\ell} \rangle,$$

$$\tilde{c}_\ell = \vec{c}_\ell - \vec{a} = \langle x_{c_\ell} - x_a, y_{c_\ell} - y_a \rangle = \langle \tilde{x}_{c_\ell}, \tilde{y}_{c_\ell} \rangle,$$

respectively. Suppose there are $m$ and $n$ lines in treatments $\boldsymbol{B}$ and $\boldsymbol{C}$**,** respectively. We denote the maximal deviations in the two dimensions as:

$$\tilde{x}_{\max} = \max\{\tilde{x}_{b_1}, \tilde{x}_{b_2}, \ldots \tilde{x}_{b_m}, \tilde{x}_{c_1}, \tilde{x}_{c_2}, \ldots \tilde{x}_{c_n}\},$$

$$\tilde{y}_{\max} = \max\{\tilde{y}_{b_1}, \tilde{y}_{b_2}, \ldots \tilde{y}_{b_m}, \tilde{y}_{c_1}, \tilde{y}_{c_2}, \ldots \tilde{y}_{c_n}\}.$$

We then express the evolved phenotypic deviations as proportions of these maximal deviations. Specifically, line $\ell$ of the $\boldsymbol{B}$ and $\boldsymbol{C}$ treatments, are expressed as:

$$\hat{b}_\ell = \langle \frac{\tilde{x}_{b_\ell}}{\tilde{x}_{\max}}, \frac{\tilde{y}_{b_\ell}}{\tilde{y}_{\max}} \rangle = \langle \hat{x}_{b_\ell}, \hat{y}_{b_\ell} \rangle,$$

$$\hat{c}_\ell = \langle \frac{\tilde{x}_{c_\ell}}{\tilde{x}_{\max}}, \frac{\tilde{y}_{c_\ell}}{\tilde{y}_{\max}} \rangle = \langle \hat{x}_{c_\ell}, \hat{y}_{c_\ell} \rangle.$$

Consider the example in Figure S5a, where there are 3 lines of treatment $\boldsymbol{B}$ and 6 lines of treatment $\boldsymbol{C}$. Generally, we see that the treatment $\boldsymbol{B}$ lines have high $\hat{x}$ values and low $\hat{y}$ values, while the treatment $\boldsymbol{C}$ lines have high $\hat{y}$ values and low $\hat{x}$ values. Of course, the high values are expected because the two sets of lines were selected for increases in those dimensions. It is the low values that are our focus here.

Figure S5b shows one statistic to summarize the tendency for relatively high gains in one dimension to correspond to relatively low gains (or even losses) in another dimension. Essentially, we draw a "summary vector" taking the average angle of all the vectors in a treatment (thick lines), and the angular distance between the two summary vectors gives a misalignment statistic that we call θ* (in pink). We note that when high gains in one dimension lead to low gains (or losses) in the other dimension, this statistic will be larger.

In Figure S6, we collect all the $\hat{x}$ values (Fig. S6a) and all the $\hat{y}$ values (Fig. S6b) of the lines from both treatments, and we plot them along the corresponding axes.

**a** **b**

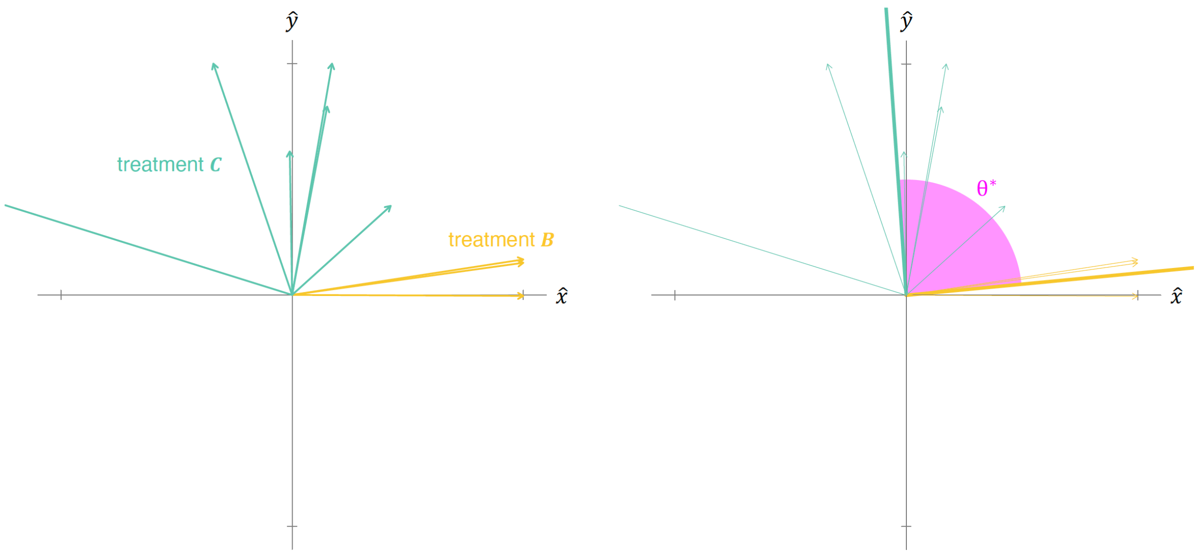


**Figure S5: (a)** The rescaled phenotypic deviations of evolved lines from two treatments (3 lines from treatment ***B*** and 6 lines from treatment ***C***) are shown as vectors, with the origin representing the common ancestral reference. The notches on the abscissa and ordinate correspond to values of 1 and -1. One of the vectors for treatment ***C*** has a very low x-value (which is allowed), and thus we do not show its arrow head. **(b)** The summary vectors (thick lines) take the average angle of all the vectors for the replicate lines in a treatment. The angular distance between the two summary vectors (θ*, in pink) is our misalignment statistic.

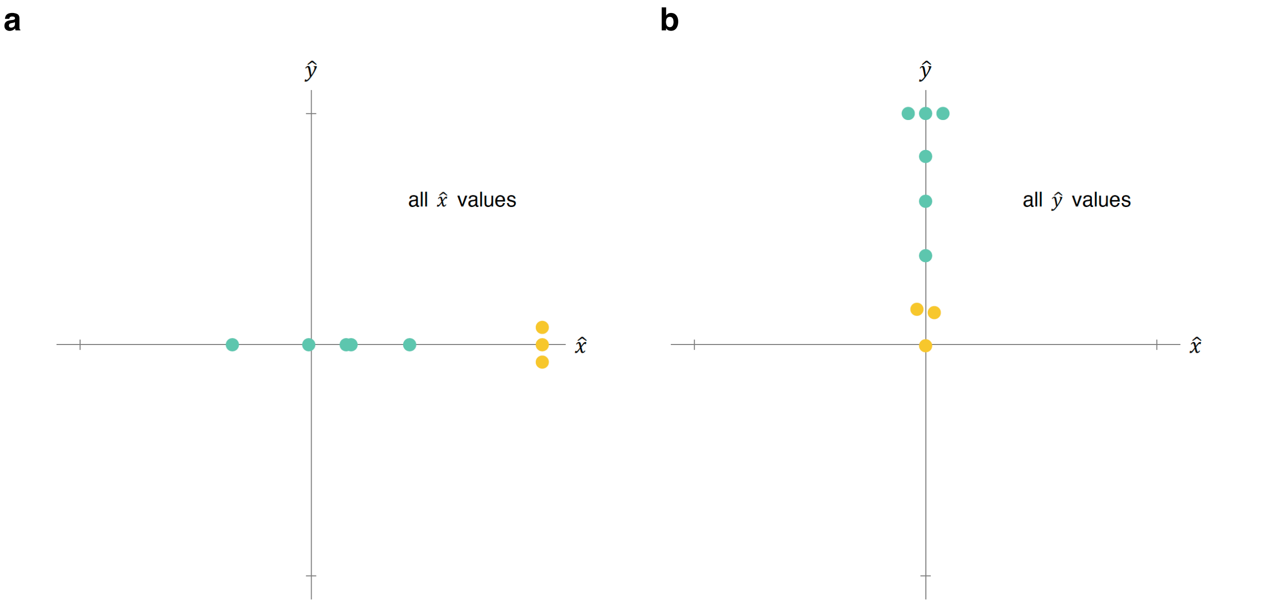


**Figure S6: (a)** Using the case illustrated in Fig. S5a, the $\hat{x}$ values are shown, with similar or equal $\hat{x}$ values separated to keep track of all values. We note that one $\hat{x}$ value (corresponding to the vector from Fig. S5a with a very low x-value) is not shown. **(b)** Using the same case, all $\hat{y}$ values are shown, again with similar or equal $\hat{y}$ values separated for visualization.

Consider the following null hypothesis: when there is not selection for the trait along the abscissa, all of the possible $\hat{x}$ values in Fig. S6a are equally likely; and when there is not selection for the trait along the ordinate, all of the possible $\hat{y}$ values in Fig. S6b are equally likely. We can then use a bootstrapping approach to gauge whether our actual statistic ($\theta^*$) is more extreme than angles that we would expect under this null hypothesis. To do so, we fix the $\hat{x}$ values of the treatment ***B*** lines and resample with replacement three $\hat{y}$ values from Fig. S6b. Similarly, we fix the $\hat{y}$ values of the treatment ***C*** lines and resample with replacement six $\hat{x}$ values from Fig. S6a. A $\theta$ statistic can then be computed on the vectors resulting from the resampling. We performed the resampling and calculation 10,000 times. Figure S7 shows 25 of these iterations, where blue indicates a value of $\theta$ smaller than our actual statistic $\theta^*$ (Fig. S5b) and red indicates a value of $\theta$ larger than $\theta^*$.

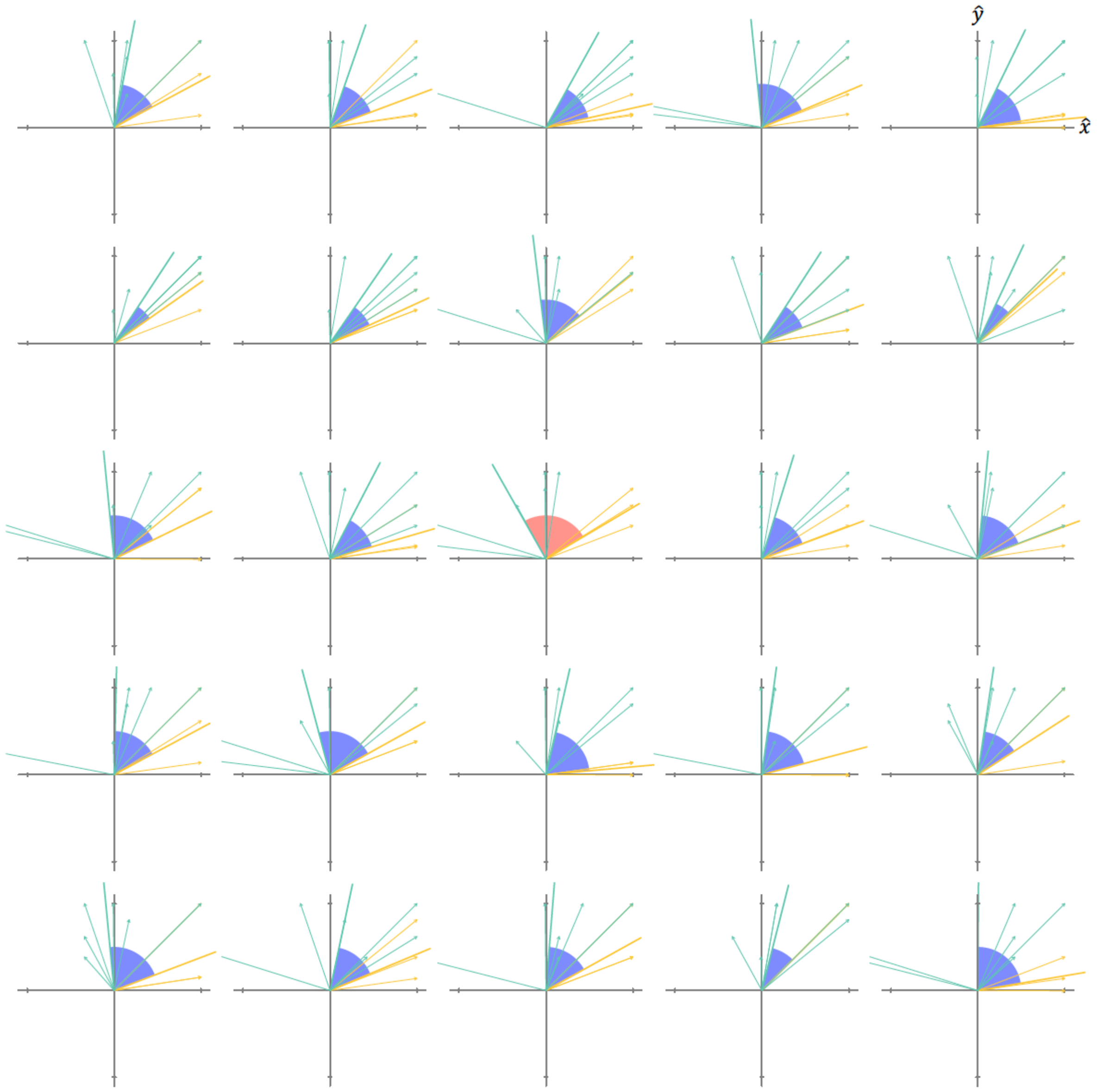


**Figure S7:** The angle between summary vectors (θ) is shown for 25 iterations of a bootstrapping scheme, in which treatment $\boldsymbol{B}$ vectors resample $\hat{y}$ values and treatment $\boldsymbol{C}$ vectors resample $\hat{x}$ values. When this resampled angle is smaller than $\theta^*$, it is colored in blue; when larger, it is colored in red.

For our case study here, we see that the $\theta$ from resampled values is more extreme than the actual statistic $\theta^*$ only 3.3% of the time (Figure S8). Thus, by convention, we would reject the null hypothesis. In this case, we would say that the responses to selection in the two treatments is significantly misaligned.

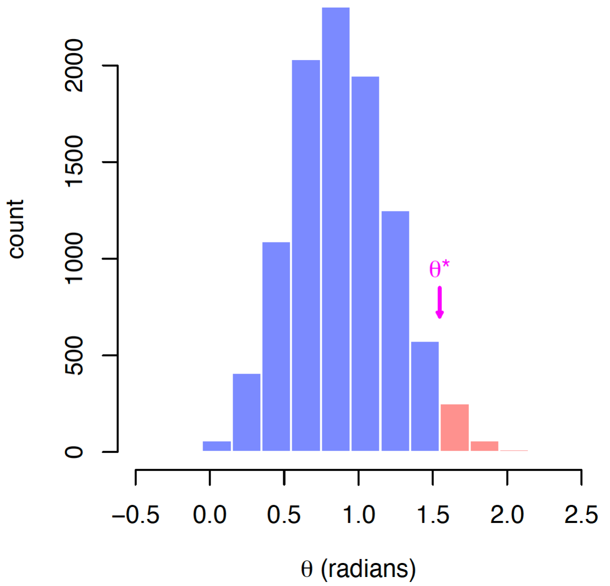


**Figure S8**: The distribution of angles resulting from our bootstrapping procedure. Only 3.3% of the angles from resampled data are more extreme than our actual misalignment statistic ($\theta^*$). The red portion of the distribution corresponds to larger angles than the actual statistic, and the blue corresponds to smaller angles.

Our case study uses the actual data from our emulsion evolution experiments. However, there are three caveats that must be made clear. First, for each emulsion ancestor (0K, 20K, and 60K) the evolved competitive fitness (abscissa) and numerical yield (ordinate) in the shared-droplet and private-droplet treatments were rescaled relative to the phenotypic values of their common ancestor. Thus, the 0K descendants were rescaled relative to the 0K ancestor, the 20K descendants relative to the 20K ancestor, and the 60K descendants relative to the 60K ancestor. Second, all of the rescaled data were pooled for our misalignment analysis (including six private-droplet lines and three shared-droplet lines). Third, we note that these data are from only part of the experiment. The only private-droplet lines that we analyzed were those exhibiting a mutation called by our first pass of genomic sequencing, and each shared-droplet descendant line was randomly picked from the three shared-droplet descendant lines for each emulsion ancestor. With these caveats in mind, the analysis is consistent with a misalignment between competitive ability and numerical yield.

**List of Strains and Supplemental Results**

| STRAIN (BK #) | LTEE SOURCE GENERATION | LTEE ID | TREATMENT (THIS STUDY) | PHENOTYPIC ASSAYS RUN? |
|---|---|---|---|---|
| **427** | 0K | REL 606 | None | Yes |
| **429** | 20K | REL 8597 | None | Yes |
| **430** | 60K | REL 11700 | None | Yes |
| **431** | 0K | NA | Private droplet | Yes |
| **432** | 0K | NA | Private droplet | No |
| **433** | 0K | NA | Private droplet | Yes |
| **437** | 20K | NA | Private droplet | Yes |
| **438** | 20K | NA | Private droplet | Yes |
| **439** | 20K | NA | Private droplet | No |
| **441** | 60K | NA | Private droplet | Yes |
| **442** | 60K | NA | Private droplet | Yes |
| **443** | 0K | NA | Shared droplet | Yes |
| **445** | 20K | NA | Shared droplet | Yes |
| **446** | 20K | NA | Shared droplet | No |
| **447** | 60K | NA | Shared droplet | No |
| **448** | 60K | NA | Shared droplet | Yes |
| **449** | 60K | NA | Shared droplet | No |

**Table S1:** Strains used and generated in this study.

| STRAIN (BK #) | LTEE GEN. | EVOL-UTION | COMPETITIVE ABILITY | Δ COMPETITIVE ABILITY | YIELD (CELLS/ML) | Δ YIELD (CELLS/ML) | CELL VOLUME ($\mu M^3$) | Δ CELL VOLUME ($\mu M^3$) |
|---|---|---|---|---|---|---|---|---|
| **431** | 0K | Private droplet | 1.03 | 0.02 | 1.82E+09 | 3.57E+08** | 0.249 | -0.013 |
| **433** | 0K | Private droplet | 1.01 | -0.001 | 1.69E+09 | 2.23E+08* | 0.246 | -0.016 |
| **437** | 20K | Private droplet | 1.02 | -0.06 | 1.74E+09 | 7.72E+08** | 0.323 | -0.276** |
| **438** | 20K | Private droplet | 1.11 | 0.03 | 1.60E+09 | 6.28E+08** | 0.371 | -0.228** |
| **441** | 60K | Private droplet | 1.01 | -0.18** | 1.70E+09 | 7.48E+08** | 0.325 | -0.304** |
| **442** | 60K | Private droplet | 1.21 | 0.02 | 1.24E+09 | 2.88E+08** | 0.529 | -0.101** |
| **443** | 0K | Shared droplet | 1.14 | 0.13* | 1.51E+09 | 5.00E+07 | 0.311 | 0.049** |
| **445** | 20K | Shared droplet | 1.25 | 0.17 | 9.68E+08 | -3.33E+06 | 0.634 | 0.036 |
| **448** | 60K | Shared droplet | 1.24 | 0.05 | 1.06E+09 | 1.15E+08* | 0.702 | 0.072 |

**Table S2:** Results of unpaired, two-tailed t-tests comparing each evolved strain with its corresponding LTEE ancestor. Significance levels are denoted by single asterisk (* $p < 0.05$) and double asterisk (** $p < 0.01$).

| STRAIN (BK #) | LTEE GEN. | GENOME POSITION | REF. | ALT. | ANNOTATION | GENE | DESCRIPTION |
|---|---|---|---|---|---|---|---|
| **431** | 0K | 558049 | G | A | G234E (GGA→GAA) | *ECB_RS02700* → | hypothetical protein |
| **431** | 0K | 1352719 | G | T | Q57K (CAA→AAA) | *ECB_RS06750* ← | peptide ABC transporter permease SapC |
| **433** | 0K | 1316697 | T | A | H229L (CAC→CTC) | ECB_RS06560 ← | bifunctional indole-3-glycerol-phosphate synthase TrpC/phosphoribosylanthranilate isomerase TrpF |
| **433** | 0K | 4417026 | C | A | intergenic (-44/+64) | *ECB_RS21720* ← / ← *ECB_RS21725* | iron-sulfur cluster repair protein YtfE/DMT family transporter |
| **433** | 0K | 4616552 | G | T | V342L (GTG→TTG) | *ECB_RS22740* → | multifunctional transcriptional regulator/nicotinamide-nucleotide adenylyltransferase/ribosylnicotinamide kinase NadR |
| **443** | 0K | 3761276 | G | T | G174C (GGT→TGT) | *ECB_RS18560* → | bifunctional GTP diphosphokinase/guanosine-3',5'-bis pyrophosphate 3'-pyrophosphohydrolase |
| **437** | 20K | 2462747 | A | C | *86Y (TAA→TAC) | *ECB_RS12210* → | phosphocarrier protein Hpr |
| **437** | 20K | 3414187 | G | A | E55K (GAA→AAA) | *ECB_RS16980* → | cAMP-activated global transcriptional regulator CRP |
| **438** | 20K | 1173797 | G | A | G454S (GGT→AGT) | *ECB_RS05820* → | PTS glucose transporter subunit IIBC |
| **441** | 60K | 2462629 | T | C | L47P (CTG→CCG) | *ECB_RS12210* → | phosphocarrier protein Hpr |
| **442** | 60K | 2009534 | G | A | R91C (CGC→TGC) | *ECB_RS10145* ← | DUF496 family protein |
| **447** | 60K | 970689 | A | ATGTA | pseudogene (829/3153 nt) | *ECB_RS04795* ← | formate acetyltransferase |
| **448** | 60K | 610497 | G | A | A32A (GCG→GCA) | *ECB_RS02935* → | enterobactin biosynthesis bifunctional isochorismatase/aryl carrier protein EntB |

**Table S3:** List of mutations that distinguish evolved strains from their corresponding ancestors.

| STRAIN (BK #) | LTEE GEN. | EVOLUTION TREATMENT | ACETATE | LACTATE | FORMATE |
|---|---|---|---|---|---|
| 427 | 0K | Emulsion ancestor | 0.45806871 | 0.00081899 | 0.15687151 |
| 429 | 20K | Emulsion ancestor | 0.45432415 | 0.08338413 | 0 |
| 430 | 60K | Emulsion ancestor | 0.40907412 | 0.11643726 | 0 |
| 431 | 0K | Private droplet | 0.47546268 | 0.00245811 | 0.3117939 |
| 433 | 0K | Private droplet | 0.53013278 | 0.00227116 | 0.29917948 |
| 437 | 20K | Private droplet | 0.0166108 | 0.16105648 | 0.0072549 |
| 438 | 20K | Private droplet | 0.12603994 | 0.16403598 | 0 |
| 441 | 60K | Private droplet | 0.01745956 | 0.08400922 | 0.00393067 |
| 442 | 60K | Private droplet | 0.192521 | 0.10040607 | 0.00042268 |
| 443 | 0K | Shared droplet | 0.47493262 | 0.00132017 | 0.23256199 |
| 445 | 20K | Shared droplet | 0.65689706 | 0.00237432 | 0.25445829 |
| 448 | 60K | Shared droplet | 0.6642624 | 0.00390774 | 0.26905594 |

**Table S4**: Standardized rates of production of three excreted metabolites per unit of glucose consumed.
